# Glutamate as a co-agonist for acid-sensing ion channels to aggravate ischemic brain damage

**DOI:** 10.1101/2024.02.17.580727

**Authors:** Ke Lai, Iva Pritišanac, Zhen-Qi Liu, Han-Wei Liu, Li-Na Gong, Ming-Xian Li, Jian-Fei Lu, Xin Qi, Tian-Le Xu, Julie Forman-Kay, Hai-Bo Shi, Lu-Yang Wang, Shan-Kai Yin

## Abstract

Glutamate is traditionally viewed as the first messenger to activate N-methyl-D-aspartate receptors (NMDARs) and downstream cell death pathways in stroke^1,2^, but unsuccessful clinical trials with NMDAR antagonists implicate the engagement of other NMDAR-independent mechanisms^3–7^. Here we show that glutamate and its structural analogs, including NMDAR antagonist L-AP5 (or APV), robustly potentiated currents mediated by acid-sensing ion channels (ASICs) which are known for driving acidosis-induced neurotoxicity in stroke^4^. Glutamate increased the proton affinity and open probability of ASICs, aggravating ischemic neurotoxicity in both *in vitro* and *in vivo* models. Site-directed mutagenesis and structure-based *in silico* molecular docking and simulations uncovered a novel glutamate binding cavity in the extracellular domain of ASIC1a. Computational drug screening of NMDAR competitive antagonist analogs identified a small molecule, LK-2, that binds to this cavity and abolishes glutamate-dependent potentiation of ASIC currents but spares NMDARs, providing strong neuroprotection efficacy comparable to that in ASIC1a or other cation ion channel knockout mouse models^4–7^. We conclude that glutamate serves as the first messenger for ASICs to exacerbate neurotoxicity, and that selective blockage of glutamate binding sites on ASICs without affecting NMDARs may be of strategic importance for developing effective stroke therapeutics devoid of the psychotic side effects of NMDAR antagonists.

---

Ischemic stroke remains the leading cause of death and disability in the world^8^. During stroke, ischemia results in a deficiency of glucose and oxygen, leading to excessive glutamate release and accumulation in the infarct area where it reaches 10-100 fold higher concentrations than normal physiological levels in rodents^1,9^. This causes overactivation of NMDARs and subsequent intracellular calcium overload, triggering cell death pathways. NMDAR-dependent excitotoxicity thus points to NMDARs as a key target for stroke treatments^2^. Indeed, numerous studies over the past decades have proven that blockage of NMDARs in neurons *in vitro* and animal models of stroke *in vivo* is highly effective in protecting neurons from injury and death ^10–13^. However, the effectiveness of NMDAR antagonists in hundreds of promising preclinical animal studies have failed to translate from bench to bedside at the stage of clinical trials^3^, raising the possibility that NMDARs are not solely accountable for glutamate-induced excitotoxicity in stroke.

Acid-sensing ion channels (ASICs), which are abundantly expressed in the central nervous system (CNS)^14,15^, have been proposed as a strong candidate for mediating glutamate-independent neurotoxicity since local acidosis occurs in ischemic areas during stroke^4,16,17^. Previous studies have suggested that NMDARs and ASICs may be coincidently activated by excessive glutamate and H^+^, respectively, and that these channels also cross talk via intracellular couplings to drive neuronal hyperexcitability and Ca^2+^ overload, aggravating neuronal death under ischemic conditions^5^. Aside from direct glutamate-gated activation of NMDARs and their downstream intracellular signaling pathways, it is unknown whether glutamate can act directly on ASICs to mediate and/or aggravate ischemic brain injury.

## Glutamate and its analogs potentiate I_ASICs_

To directly investigate whether the function of ASICs is regulated by glutamate, we performed patch-clamp experiments to record acid-evoked currents (I_ASICs_) from CHO cells expressing ASIC1a channels and primary cultured cortical neurons. We found that glutamate, but not glutamic acid monosodium, potently potentiated I_ASICs_ evoked by solutions with pH values ranging from 6.55 to 7.4 (i.e. 0.4 to 2.8 × 10^-7^ mol/L proton concentrations). The maximal potentiation was achieved at ∼pH 6.85 and the potentiation diminished at pH<6.5 (Fig. 1a and Extended Data Fig. 1a-f). This potentiation was concentration-dependent with an EC_50_ of 473.3 μM for glutamate, showing typical features of glutamate being a positive allosteric modulator (Fig. 1b). Ca^2+^ is thought to block ASIC1a^18^, raising the possibility that glutamate may attenuate the action of Ca^2+^, unmasking the inhibition of I_ASICs_. To this end, we generated an ASIC1a mutant that ablated the Ca^2+^ blocking site (*h*ASIC1a^E427G/D434C^). We found that the amplitudes of *h*ASIC1a^E427G/D434C^ currents were decreased compared to that of wildtype currents, likely due to the impaired surface expression of functional mutants and/or its conductance. Nevertheless, glutamate still potentiated *h*ASIC1a^E427G/D434C^ currents at pH above 6.65, consistent with the results in wildtypes (Extended Data Fig. 1g). Because ASICs can be present either as homomeric ASIC1a or heteromeric ASIC1a/2a and ASIC1a/2b channels in native central neurons^19^, we tested the effect of glutamate on these heteromeric channels and noted that the potentiation by glutamate was preserved (Fig. 1c). These results indicated that ASIC1a mediated the effect of glutamate on ASICs, which was largely conserved, albeit with subtle differences, regardless of their subunit composition. Thus, glutamate appears to act as a co-agonist to allosterically potentiate ASICs, reminiscent of glycine as a co-agonist for NMDARs.

**Figure 1.**
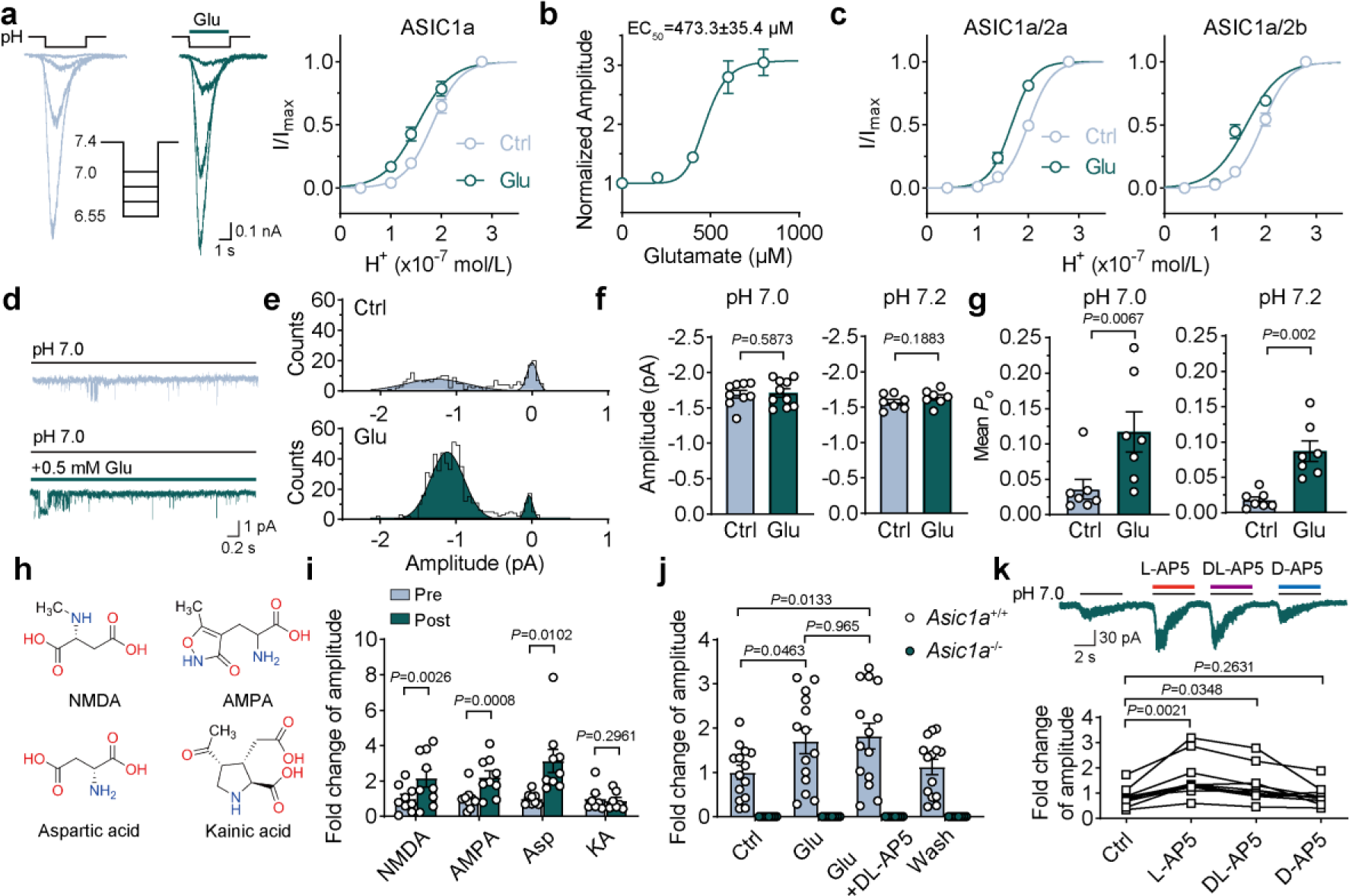
Glutamate and its structural analogs, including AP5, robustly potentiate ASIC currents. **a**, Left: ASIC currents (I_ASICs_) evoked by stepwise changes in solution with pH value from 7.4 to 7.0, 6.85, 6.7 and 6.55 in an ASIC1a transfected CHO cell in the absence and presence of 500 μM glutamate. Right: pH-response curves for ASIC1a currents with and without glutamate were plotted against the H^+^ concentration. *n*=14 cells. **b**, Dose-response curve for glutamate to potentiate I_ASICs_. *n*=14 ASIC1a transfected CHO cells. **c**, Dose-response curves for ASIC1a/2a and ASIC1a/2b currents with and without glutamate (500 μM). *n*=13 and 14 cells, respectively. **d**, Outside-out patch recordings of ASIC single-channel currents evoked by pH 7.0 before and after addition of 500 μM glutamate. Currents were recorded at -60 mV. Data were filtered at 1 kHz. **e**, All points amplitude histogram of ASIC single-channel currents was illustrated from (**d**), curves were fitted by double Gaussian components. Bin=0.05 pA. **f**,**g**, Quantification of amplitude and mean open probability (*P*_o_) of ASIC1a unitary currents evoked in the presence and absence of 500 μM glutamate at pH 7.0 and 7.2. *n*=7-9 patches. **h**, Chemical structure of glutamate analogs. **i**, Quantification of I_ASICs_ recorded before and after 500 μM NMDA, AMPA, aspartic acid (Asp) and kainic acid (KA) treatment at pH 7.0. **j**, Quantification of I_ASICs_ recorded from primary cultured cortical neurons from *Asic1a*^+/+^ (*n*=14 cells) and *Asic1a*^-/-^ mice (*n*=8 cells) at pH 7.0. 500 μM glutamate and 200 μM DL-AP5 were used. **k**, Representative traces (superimposed) and summary data showing the effect of L-, DL- and D-isomers of AP5 (400 μM) on ASIC1a at pH 7.0 (*n*=11 cells). Data are mean±s.e.m.; two-tailed paired Student’s *t*-test (**f**, **g**, **i**); two-way analysis of variance (ANOVA) with Tukey post hoc correction for multiple comparisons (**j**); oneway analysis of variance (ANOVA) with Dunnett post hoc correction for multiple comparisons (**k**). *P* values are indicated.

To investigate the biophysical basis of potentiation of I_ASICs_ by glutamate, we performed single-channel recordings in outside-out patches from ASIC1a transfected CHO cells. We found that glutamate increased the open probability (*P*_o_) of ASIC1a at pH 7.0 and even a milder pH 7.2, without affecting the amplitude of ASIC1a unitary currents at both pH 7.0 and 7.2 (Fig. 1d-g). In addition, the current-voltage relationship before and after bath application of glutamate showed the same slope values (conductance: 17.3±0.9 pS; glutamate: 18.4±0.6 pS) and reversal potential (control: 33.15 mV; glutamate: 28.56 mV) (Extended Data Fig. 1h, i), indicating that glutamate-induced potentiation of I_ASICs_ was solely accounted for by an increase in *P*_o_ while single-channel conductance and ion selectivity of ASICs were not affected.

To further examine the potentiation of I_ASICs_ by glutamate in ASIC1a transfected CHO cells, we tested other structural analogs, including *N*-methyl-*D*-aspartic acid (NMDA), amino-3-hydroxy-5-methylisoxazole-4-propionic acid (AMPA), aspartic acid (Asp) and kainic acid (KA). Similar to glutamate, NMDA, AMPA and Asp all potentiated I_ASICs_, while KA had no effect (Fig. 1h, i). We replicated these results in primary cultured cortical neurons and confirmed that glutamate significantly potentiated I_ASICs_ at pH 7.0 in neurons from wild-type mice (*Asic1a*^+/+^). This potentiation was completely absent in those from ASIC1a knockout mice (*Asic1a*^-/-^) (Fig. 1j). These results prompted us to investigate whether this potentiation could be blocked by the classical NMDAR competitive blocker 2-amino-5-phosphonovaleric acid (AP5 or APV). We first tested the effect of three isomers (L-, DL- and D-) of AP5 on I_ASICs_. None of the AP5 isomers (200 μM) abolished glutamate-dependent enhancement of I_ASICs_ in CHO cells (Extended Data Fig. 2a, b). Surprisingly, we observed that L- and DL-but not D-isomer of AP5 enhanced I_ASICs_ (Fig. 1k). Like the effect of glutamate, DL-AP5 increased the *P*_o_ of ASIC1a at pH 7.0 and even at a milder pH 7.2, demonstrating the chiral effect of AP5 on ASIC1a resides in its L-isomer (Extended Data Fig. 2 c-f).

Taken together, these observations suggested that ASIC1a exhibits glutamate-dependent positive modulation of I_ASICs,_ and that glutamate and its structural analogs share a common binding site most likely on the extracellular domain of ASICs to allosterically potentiate I_ASICs_.

## Glutamate induces NMDAR-independent cell death

Excessive glutamate release and acidosis are believed to cause neuronal death by activating both NMDARs and ASICs in ischemic brain injury^4,20^. Our observations led us to hypothesize that glutamate may cause NMDAR-independent cell death by directly acting on ASICs to drive Ca^2+^ overload. To this end, we performed calcium imaging in cultured cortical neurons from *Asic1a*^+/+^ and *Asic1a*^-/-^ mice in the presence of AMPA receptor blocker NBQX (1,2,3,4-tetrahydro-6-nitro-2,3-dioxo-benzo[f] quinoxaline-7-sulfonamide), NMDAR pore blocker MK801 and voltage-gated calcium channel blocker CdCl_2_ (Extended Data Fig.3a). In *Asic1a*^+/+^ cortical neurons, we found application of pH 7.0 solution alone led to slow elevation of intracellular Ca^2+^ ([Ca^2+^]_i_) which was robustly potentiated by co-application of glutamate, consistent with the idea that glutamate can promote Ca^2+^ overload independent of NMDARs and other potential sources of Ca^2+^ influx. This was further reinforced by the same experiments with *Asic1a*^-/-^ neurons, in which glutamate showed no significant effects on [Ca^2+^]_i_ elevation at pH 7.0 (Fig. 2a, b).

**Figure 2.**
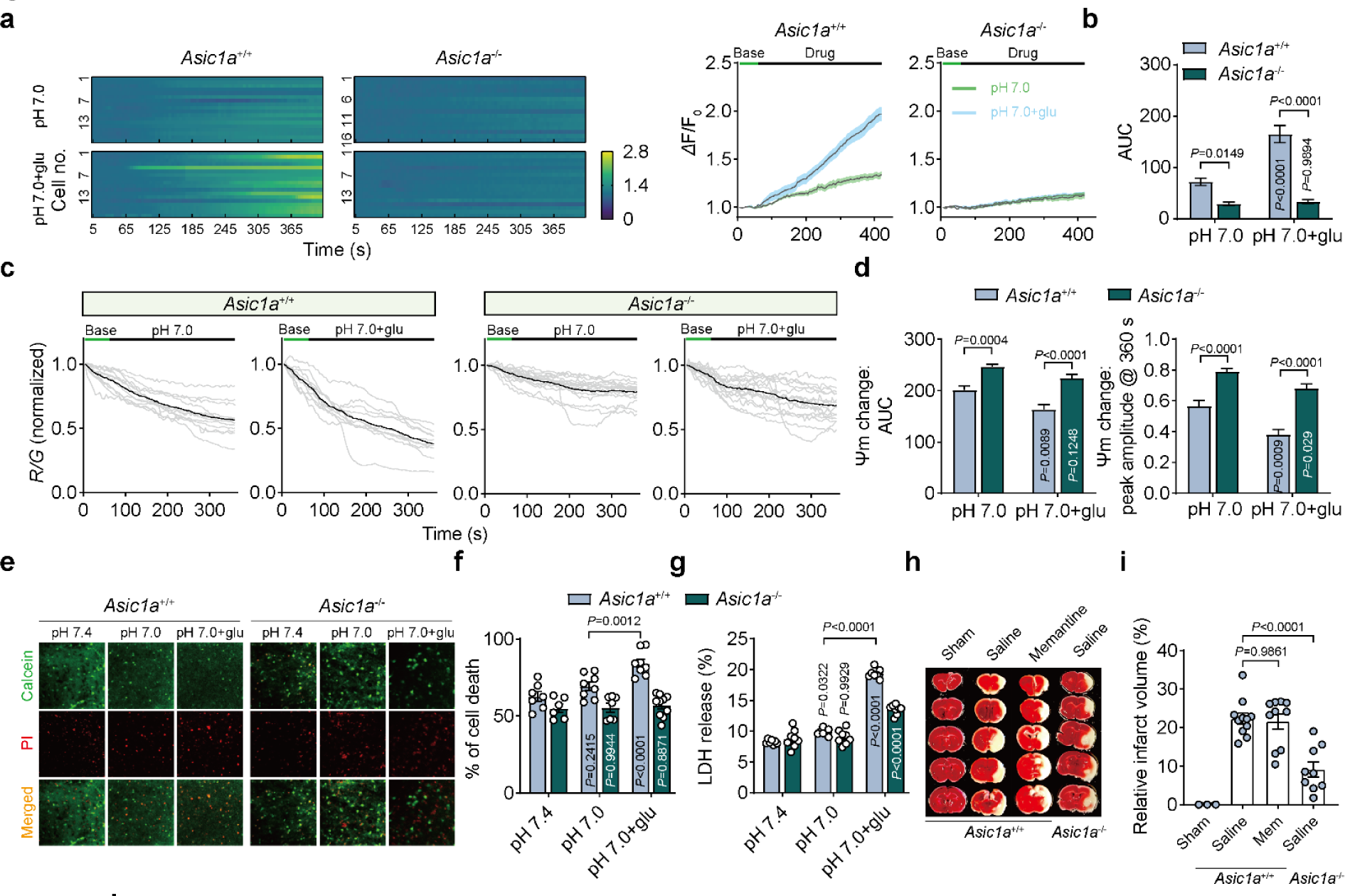
Glutamate aggravates NMDAR-independent neurotoxicity *in vitro* and *in vivo*. **a**, **b**, Calcium changes of primary cultured cortical neurons imaged from *Asic1a*^+/+^ (*n*=18 and 18 cells for each group) and *Asic1a*^-/-^ (*n*=16 and 18 cells for each group) mice with and without addition of 500 μM glutamate to pH 7.0 solution. Left (**a**), fluorescence changes of individual cells; right (**a**), average of fluorescence change over time; (**b**), area under curve (AUC) quantified from right (**a**). **c**, Fluorescence changes in response to different treatments imaged from *Asic1a*^+/+^ (*n*=12 and 10 cells for each group) and *Asic1a*^-/-^ (*n*=16 and 15 cells for each group) cultured cortical neurons. The ratio of red and green fluorescence density was normalized to their initial value. Gray lines represented responses of individual cells while black line was the mean of all cells. **d**, Summary data for the AUC (left panel) and peak response amplitude (right panel) of mitochondrial membrane potential (Ψm) changes during treatment. **e**,**f**, Representative images and summary data showing the percentage of cell death by calcein-PI staining of cortex (visual and auditory area) slices from wildtype (*n*=7, 8 and 8 slices for each group) and *Asic1a*^-/-^ (*n*=6, 7 and 10 slices for each group) mice under different conditions. Calcein (green), live cells; PI (red), dead cells. **g**, Quantification data showing LDH release of brain slices from *Asic1a*^+/+^ (*n*=7 slices for each group) and *Asic1a*^-/-^ (*n*=8 slices for each group) mice under different conditions. **h**, Images of brain slices (thickness, 1 mm) from MCAO *Asic1a*^+/+^ and *Asic1a*^-/-^ mice (sham, the same surgery process as the MCAO group except for no occlusion of the middle cerebral artery). **i**, Quantification of infarct volume after MCAO from mice injected with physiological saline or NMDAR blocker, memantine (20 mg/kg, i.p.). *n*=3, 11, 10 and 9 mice for each group. Data are mean±s.e.m.; two-way ANOVA with Tukey post hoc correction for multiple comparisons (**b**, **d**, **f**, **g**); one-way ANOVA with Tukey post hoc correction (**i**). *P* values are indicated.

Mitochondrial dysfunction is a hallmark of excitotoxicity and an early event *en route* to neuronal death^21–23^. We next tested the mitochondrial membrane potential (Ψm) by a JC-1 (5,5′,6,6′-tetrachloro-1,1′,3,3′-tetraethyl-imidacarbocyanine iodide) kit to assess its function under different conditions (Extended Data Fig.3b). Glutamate caused a rapid drop of Ψm in *Asic1a*^+/+^ neurons at pH 7.0. By contrast, *Asic1a*^-/-^ neurons did not show any significant decrease in Ψm following co-application of glutamate and pH 7.0 (Extended Data Fig.3c and Fig. 2c, d).

To address whether glutamate causes cell death through its binding to ASIC1a, we subjected freshly isolated brain slices from *Asic1a*^+/+^ and *Asic1a*^-/-^ mice to calcein-PI (propidium iodide) staining of dead cells and lactate dehydrogenase (LDH) release assay as a subjective readout of neuronal injury. Given that the cortex is clinically the main loci of injury in ischemic stroke, we focused on cell death in the visual and auditory cortex (Extended Data Fig.4a). We found that glutamate markedly increased the percentage of cell death in *Asic1a*^+/+^ slices, whilst in contrast, *Asic1a*^-/-^ slices had decreased cell death and seemed resistant to glutamate insults (Fig. 2e,f). ASICs are expressed throughout the whole brain, so we also quantified the percentage of cell death in other brain areas, e.g. media vestibular nucleus (MVN) in the brainstem (Extended Data Fig.4a). We found that *Asic1a*^-/-^ MVN slices displayed no significant increase in cell death following glutamate treatment compared to the baseline condition (pH 7.4) (Extended Data Fig.4b, c). Compared to slices treated at pH 7.4, slight acidification of extracellular fluid alone had very little effect, but addition of glutamate at pH 7.0 induced a significant increase in cell death and LDH release in *Asic1a*^+/+^ slices. Again, these changes were largely attenuated in *Asic1a*^-/-^ slices (Fig. 2g). Collectively, these three sets of experiments *in vitro* demonstrated that glutamate plays a vital role in amplifying the level of [Ca^2+^]_i_, mitochondrial dysfunction and cell injury/death even in a slightly acidic condition (e.g. pH 7.0), lending support to the idea that glutamate works through ASIC1a to cause NMDAR-independent Ca^2+^ overload and neurotoxicity.

To further address the hypothesis that ASICs mediate NMDAR-independent brain damage *in vivo*, we established an ischemic stroke mouse model by performing transient middle cerebral artery occlusion (MCAO) for 30 min to induce excessive glutamate release and acidosis during ischemia and reperfusion. The infarct volume was quantified by post-hoc tissue staining with 2,3,5-triphenyltetrazolium chloride (TTC) 24 hrs later (Extended Data Fig.4d). MCAO reliably induced diminishment of relative cerebral blood flow (rCBF) and cerebral infarction during surgery, but administration of memantine (NMDAR open channel blocker, 20 mg/kg, i.p.) failed to show any protection. In contrast, significantly smaller infarct volumes were noted in *Asic1a*^-/-^ mice than in *Asic1a*^+/+^ mice while the rCBF during MCAO between the two genotype mice showed no difference (Extended Data Fig.4e,f and Fig. 2h,i). Together, these data indicated that ischemic brain damage in an MCAO mouse model was NMDAR-independent and largely ASIC1adependent.

## Glutamate binds to an extracellular pocket of ASIC1a

Given our findings that glutamate and its chemical analogs acted directly on ASIC1a to enhance I_ASICs_, we hypothesized that there might be a binding pocket for glutamate and its analogs on ASIC1a to enable their role as co-agonists. To gain insights into the molecular basis for glutamate binding to ASIC1a, in-depth analyses of the properties and conservation of the accessible surface area in the ASIC1a structure as well as computational docking were performed to generate binding site predictions for glutamate. We focused our analyses on the trimeric extracellular domain of the available chicken ASIC1a (*c*ASIC1a) structure (PDB: 5WKU) in a homomeric subunit channel assembly^24^. At the onset of this work, only a low resolution cryo-electron structure of the human protein was available^25^, so we instead performed computational simulations based on a higher resolution structure of *c*ASIC1a which shares significant conservation with human ASIC1a sequences (89.5%, Extended Data Fig. 5). We identified six strong candidate sites around amino acid residues R161, K379, K383, Q226, K391 and K387 (Fig. 3a). In addition, we performed molecular dynamics simulations to provide information on the energetics and spatial constraints to establish the rank-order for potential glutamate binding sites (Extended Data Table.1). We remapped these putative sites back onto the human ASIC1a (*h*ASIC1a) sequence, with one amino acid sequence shift (i.e., R160, K380, K384, Q225, K392 and K388) akin to those in *c*AISC1a.

**Figure 3.**
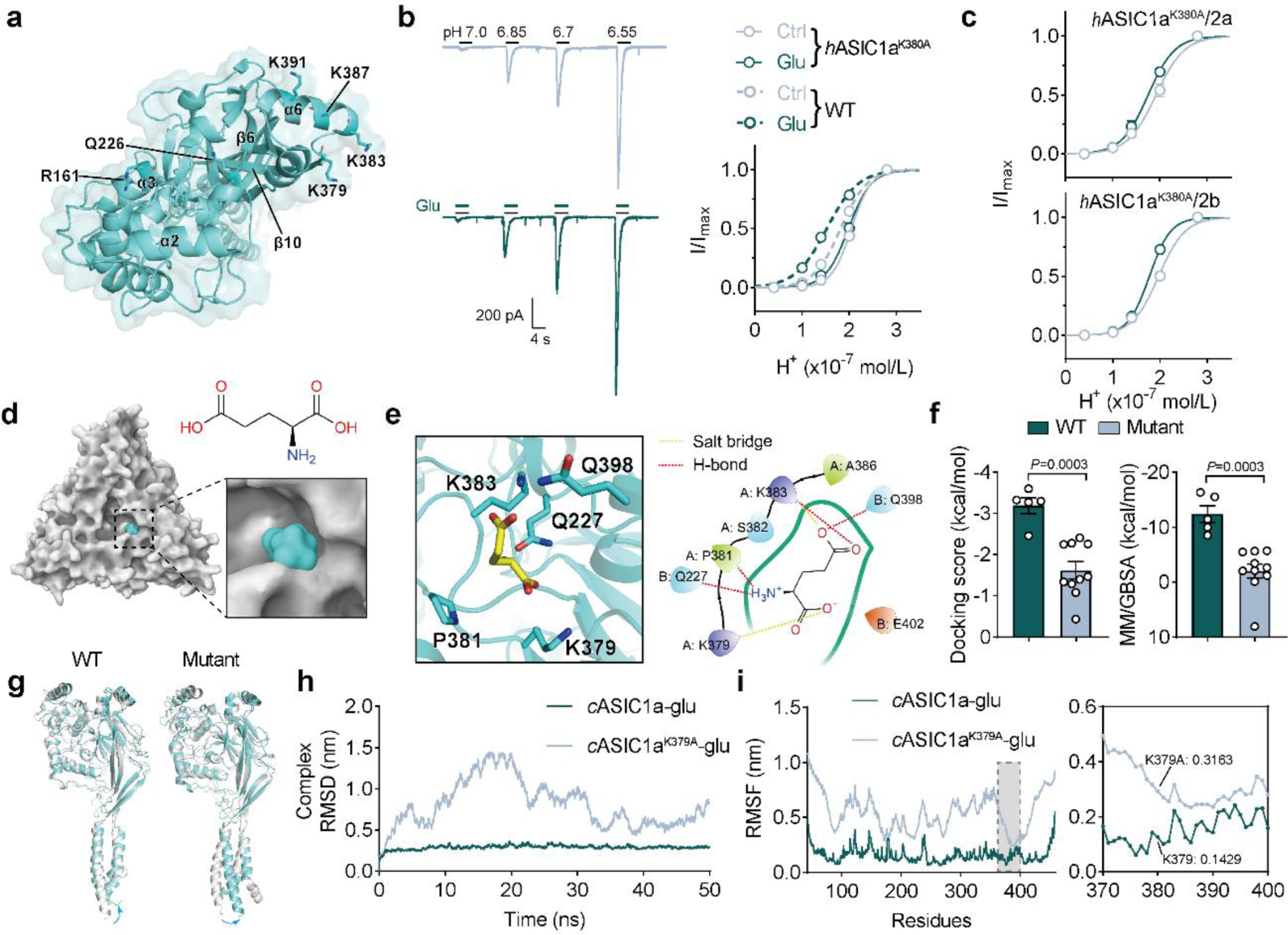
Structure-based determination of glutamate binding pocket in the extracellular domain of ASIC1a. **a**, A top view of chicken ASIC1a (PBD: 5WKU). Six highest scoring binding residues of glutamate by initial docking calculation are shown. For clarity, only chain A is shown here. **b**, Example traces of I_ASICs_ activated by different pH solutions in a human ASIC1a^K380A^ transfected CHO cell in the presence and absence of 500 μM glutamate (left panel). Dose-response curves (solid lines) were contrasted to those from homomeric ASIC1a channels as in figure 1a (dashed lines). *n*=12 cells for ASIC1a^K380A^. **c**, Dose-response curve showing I_ASICs_ activated by the same protocol in (**b**) from *h*ASIC1a^K380A^/2a, *h*ASIC1a^K380A^/2b transfected CHO cells. *n*=10 cells for each group. **d**, Top view of glutamate-bound *c*ASIC1a. Close-up view of the glutamate-binding pockets are shown in the right lower inset. Chemical structure of glutamate is shown in the right upper inset. The surface of *c*ASIC1a and glutamate are colored white and cyan, respectively. **e**, Three-dimension and two-dimension images showing glutamate bounded pocket. The interactions between glutamate and surround residues are shown as yellow (salt bridge) and red (hydrogen-bond) dash lines. **f**, Summary data by computational calculation showing MM/GBSA and docking score of glutamate binding to *c*ASIC1a. Mutant indicates residue K379 mutated to alanine. *n*=5 and 10 poses, respectively. **g**, Conformational changes of wildtype and mutant *c*ASIC1a before (white) and after (cyan) 50 ns MD simulation. Structures were aligned. For clarity, only chain A was shown here. Arrows indicated the direction of conformational change. **h**, Structural stability of glutamate-*c*ASIC1a complex in wildtype and mutant conformations were measured as the RMSD (unit: nm) over a 50-ns time course. **i**, Structural stability of glutamate-*c*ASIC1a complex in wildtype and mutant conformations were measured as the RMSF (unit: nm) throughout chain A residues. Zoom-in views of RMSF from residues 370 to 400 are shown in the right panel. Data are mean±s.e.m. Two-tailed unpaired *t*-test (**f**). *P* values are indicated.

We next introduced mutations by site-directed mutagenesis to replace these six plausible interface residues with alanine (A) or leucine (L) in human ASIC1a and analyzed the functional consequences in these mutants through patch-clamp experiments. We found that currents in cells from the *h*ASIC K380A mutant and even the ASIC K378A mutant displayed significantly diminished sensitivity to glutamate at working pH (6.55-7.0) compared to wildtype (Fig. 3b and Extended Data Fig. 6a). In contrast, no changes in the magnitude of potentiation were observed in *h*ASIC1a K384A, Q225L and K388A mutants as compared to wildtype. Glutamate was also unable to enhance currents of *h*ASIC1a K392A and R160A mutants (Extended Data Fig.6b). The 3D structural proximity of K380 and K392 indicated these two residues as a part of the same binding pocket. The R160 mutant was not pursued further because the R160A mutant generated very little currents with markedly altered kinetics of I_ASICs_ by *h*ASIC1a, making it difficult to ascertain whether this mutation perturbed glutamate and/or proton binding or channel gating. Finally, potentiation of I_ASICs_ by glutamate in CHO cells co-expressing the *h*ASIC1a^K380A^ mutant subunit and ASIC2a or 2b subunit was significantly reduced (Fig. 3c). These data indicated that K380 in *h*ASIC1a is required for glutamate binding, likely representing a key residue in the pocket to enable its role as a co-agonist. Our patch-clamp experiments indicated that the K380A mutant did not have adverse effect on the stability of the protein as its currents appeared to be comparable. In addition, predictions of the protein stability with the K380A mutation suggest no impacts on either trimer or monomer stability^26^ (Extended Data Table 2).

To gain detailed insights into glutamate interactions within the proposed pocket around K380 of *h*ASIC1a, we performed independent molecular docking analyses using Maestro software to place glutamate near the chain A of *c*ASIC1a^K379^ at the outer vestibule of the ion permeation pathway (Fig. 3d). In Figure 3e, the optimal docking pose to the putative *c*ASIC1a binding site for glutamate is shown with optimized side-chain positions of five interacting residues in the vicinity. Glutamate appears to form four specific hydrogen bonds (H-bonds) to Q227, P381, K383 and Q398, two salt bridges to K379 and K383 (Fig. 3e), giving a very stable glutamate-*c*ASIC1a complex.

To determine whether K379 of *c*ASIC1a was indeed the core binding site for glutamate, we analyzed the molecular mechanics/generalized born surface area (MM/GBSA), binding free energy and docking score^27^ of glutamate-*c*ASIC1a^K379^ and glutamate-*c*ASIC1a^K379A^ complex models. We found that both parameters for the wildtype were significantly more negative than those of *c*ASIC1a^K379A^-glutamate complex (Fig. 3f), implicating a destabilization of the complex by K379 mutation. Given that glutamate is a natural agonist for NMDARs, therefore, we also constructed a docking model of the glutamate-NMDAR complex to identify any differences when compared to our glutamate-*c*ASIC1a complex model (Extended Data Fig.7a). Interestingly, we found no difference in docking score between glutamate-NMDAR and -*c*ASIC1a^K379^ (wildtype) complexes, whereas the MM/GBSA of glutamate-NMDAR was higher than that of the ASIC1a^K379^–glutamate complex (Extended Data Fig.7b, c), providing an explanation for glutamate binding NMDARs with a higher affinity than ASIC1a.

We next performed a 50-ns long molecular dynamics (MD) simulations on glutamate-*c*ASIC1a^K379^ (wildtype) and glutamate-*c*ASIC1a^K379A^ mutant complexes to detect any conformational changes induced by ligand binding and assess the structural stability of the protein-ligand association (Extended Data Fig.7d). Structural stability of the systems and deviation from the starting atomic positions were evaluated using the root mean square deviation (RMSD) and root mean square fluctuation (RMSF) values extracted from MD simulation trajectory during these 50 ns bouts. Interestingly, the K379A mutant appeared to induce more substantial structural rearrangements of the glutamate-bound ASIC1a compared to the wildtype model (Fig. 3g), as reflected by larger RMSD fluctuations in the glutamate-*c*ASIC1a^K379A^ model while the glutamate-*c*ASIC1a model maintained a relatively stable structure during the 50-ns simulations (Fig. 3h and Extended Data Fig.7e). In addition, the mutant complex displayed strikingly larger fluctuations of RMSF and lower binding energy for glutamate (Fig. 3i and Extended Data Fig.7f,g). Taken together, our simulations demonstrated that K379 of *c*ASIC1a (i.e., homologous site K380 of *h*ASIC1a) is the crucial amino acid residue to render the optimal binding site for glutamate to potentiate the activity of ASIC1a channels.

## Small molecules provide neuroprotection

Our findings with structure-based ligand-receptor interactions raised the possibility of developing therapeutics to alleviate neuronal injury by targeting the glutamate binding site on ASIC1a. In this regard, we tested a series of candidate chemicals to block the potentiation of I_ASICs_ by glutamate. Both 6-cyano-7-nitroquinoxaline-2,3-dione (CNQX, a competitive AMPA/kainate glutamate receptor antagonist) and memantine were unable to pharmacologically affect glutamate-dependent potentiation of I_ASICs_ because their chemical structures are distinct from the backbone of glutamate (Extended Data Fig.8a-e). CGP39551 and 4-(3-phosphonopropyl)-piperazine-2-carboxylic acid (CPP), two AP5 analogs, were also unable to affect I_ASICs_ or block the potentiation of I_ASICs_ by glutamate (Extended Data Fig.8f-k). L-Aminoadipic acid (L-AA) did not block I_ASICs_, but strikingly abolished the potentiation of I_ASICs_ by glutamate (Extended Data Fig.8l-n). However, L-AA may exert agonist activity for NMDAR^28,29^ and some metabotropic glutamate receptors (mGluRs)^30,31^, making it unsuitable for ischemic stroke therapy.

Among many tested chemicals, we found that CGS19755 (also known as *Selfotel*), a rigid analog of AP5 with water solubility and blood-brain barrier (BBB) permeability, significantly attenuated glutamate-induced potentiation of I_ASICs_ in CHO cells without affecting I_ASICs_ itself (Extended Data Fig.9a-c). Similar results were also seen in cultured cortical neurons (Extended Data Fig.9d). CGS19755 reduced glutamate-induced potentiation of I_ASICs_ in a dose-dependent manner in CHO cells with a half-maximal inhibition concentration (IC_50_) of 7.689±2.603 μM (Extended Data Fig.9e). Computational calculation of molecular docking showed that the affinity of CGS19755 binding to ASIC1a^K379^ pocket was much stronger than that of glutamate (best pose of docking scores, kcal/mol: CGS19755 to ASIC1a: -6.05; glutamate to ASIC1a: -3.605) (Extended Data Fig.9f and Extended Data Table.3). Therefore, these data support the idea that CGS19755 might be a potent competitive blocker for the glutamate binding site on both ASICs and NMDARs, providing dual neuroprotection by abolishing glutamate-induced potentiation of I_ASICs_ and blocking NMDARs.

We further performed cell injury experiments in *vitro* and in *vivo*. We found that even high concentrations (e.g. 100∼1000 μM) of CGS19755 were well tolerated by the neurons and produced no signs of cell injury (Extended Data Fig.10a). In Ca^2+^ imaging experiments, the [Ca^2+^]_i_ rise evoked by glutamate application and acidosis (i.e. pH 7.0) was largely attenuated by CGS19755 (Extended Data Fig.10b-d). In parallel, measurement of the Ψm by a JC-1 kit revealed that CGS19755 virtually eliminated glutamate/acidosis-induced mitochondrial dysfunction in cortical neurons (Extended Data Fig.10e-g). Together, these findings suggested that CGS19755, by competitively blocking glutamate binding sites on ASICs and NMDARs, could attenuate [Ca^2+^]_i_ overload and mitochondrial dysfunction of neurons induced by glutamate and acidosis, potentially providing neuroprotection against NMDAR-independent and -dependent brain injury in stroke.

Despite being a highly neuroprotective both in *vitro* and in *vivo* ischemic models, clinical trial with CGS19755 on stroke was terminated due to severe side effects including psychosis as a result of blocking NMDARs^32^. NMDARs are instrumental for brain function in physiological conditions and even have pro-survival functions during stroke recovery^33^, so unfortunately, inhibition of NMDAR by its high efficacy antagonists such as CGS19755 is not viable approach for stroke therapy^34,35^. Therefore, we screened for small molecules that could largely block glutamate-enhanced ASIC activity but produce minimal effects on NMDAR activity. To this end, we carried out a CGS19755 structure-based computational drug screen, which revealed two candidate compounds. LK-1 and LK-2 were selected based on molecular docking scores (best pose of docking scores, kcal/mol: LK-1 to ASIC1a: -6.371, LK-1 to NMDAR: -4.32; LK-2 to ASIC1a: -7.039, LK-2 to NMDAR: -4.686), as well as electrophysiological and biological assays. They are presented here as prototypes of a new class of neuroprotective small molecules (Extended Data Fig.11a and Fig. 4a,b). To verify that LK-1 and LK-2 pharmacologically function with ASIC1a and NMDAR, we first co-applied glutamate and two compounds to assess their ability to disrupt the complex of glutamate and ASIC1a. LK-1 and LK-2, indeed, reduced glutamate potentiated I_ASICs_ in a dose-dependent manner (Extended Data Fig.11b and Fig. 4c). More importantly, neither compound blocked basal currents of ASIC1a (Extended Data Fig.11c). Next, we investigated the possible effects of LK-1 and LK-2 on NMDARs. In contrast to the NMDAR antagonist CGS19755 that completely blocked NMDAR currents, LK-1 and LK-2 were much less effective in attenuating NMDAR currents in acute isolated cortical neurons and CHO cells that express recombinant NMDARs (Extended Data Fig.11d and Fig. 4d). We found that NMDARs comprised of NR1 (GluN1) plus NR2A (GluN2A) subunits were more sensitive to LK-1 and LK-2 than NR1 plus NR2B (GluN2B) subunits (Fig. 4d). Moreover, LK-2 was found to be a weaker antagonist for NMDAR than LK-1 (Fig. 4d), being two orders of magnitude higher in its IC_50_. This highlighted its potential to preferentially block the glutamate binding site on ASICs over the site on NMDARs. The pharmacological effects of LK-1 and LK-2 on ASIC1a and NMDARs were in line with our docking results *in silico* that LK-1 and LK-2 both displayed higher affinity for ASIC1a than NMDAR, but LK-2 is more likely to spare NMDARs over LK-1. Indeed, in cell death and LDH release assays under the acid condition with glutamate treatment in *vitro*, LK-2 remarkably reduced cell death and LDH release to an extent close to CGS19755, indicating glutamate-enhanced ASIC activity rather than NMDAR overactivation was the main contributor to cell injury in stroke conditions (Fig. 4e-g).

**Figure 4.**
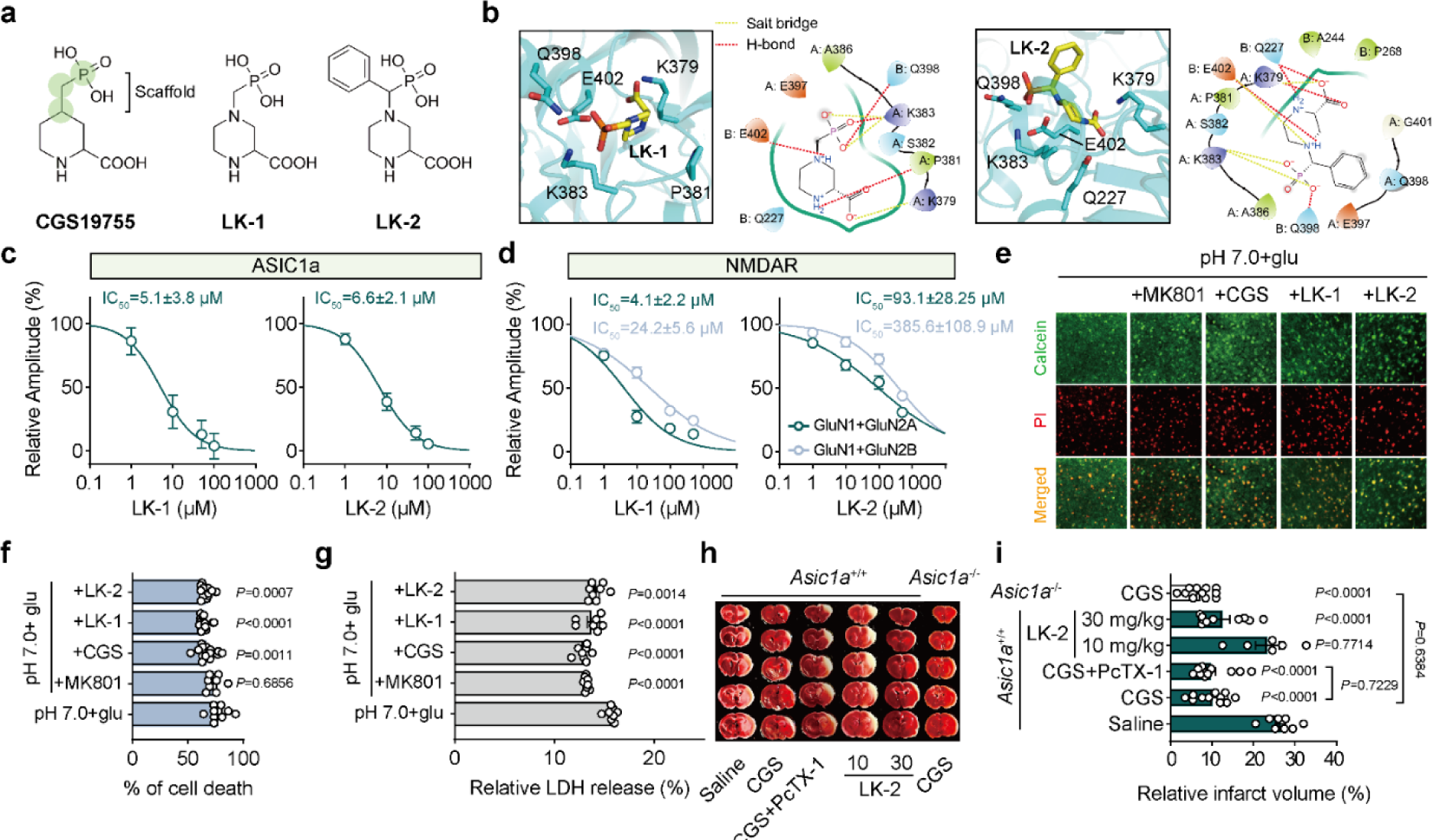
Targeting glutamate binding site on ASICs with novel compounds is protective against neurotoxicity. **a**, Chemical structures of CGS19755, LK-1 and LK-2. Green coded circles represent the scaffold for virtual screening. **b**, Three-dimension and two-dimension images showing LK-1 and LK-2 bounded pocket. The interactions between ligand and surround residues are shown as yellow (salt bridge) and red (hydrogen-bond) dash lines. **c**, Dose-response curves showing glutamate-dependent potentiation of I_ASICs_ was inhibited by LK-1 and LK-2 at pH 7.0. **d**, Dose-response curves showing LK-1 and LK-2 inhibited NMDAR currents evoked by 100 μM NMDA and 10 μM glycine. NR1 (GluN1) plus NR2A (GluN2A) or NR2B (GluN2B) subunits were co-expressed in CHO cells. **e**,**f**, Representative images and summary data showing percentage of cell death by calcein-PI staining of cortex (visual and auditory area) slices from wildtype (*n*=7, 8, 16, 14 and 13 slices for each group) mice under different conditions. Calcein (green), live cells; PI (red), dead cells. **g**, Quantification data showing LDH release of brain slices from wildtype (*n*=8, 7, 7, 7 and 8 slices for each group) mice under different conditions. 200 μM glutamate, 1 μM MK801, 100 μM CGS19755, 100 μM LK-1 and 100 μM LK-2 were applied for calcein-PI staining and LDH release assay, respectively. **h**,**i**, Images of brain slices (thickness, 1 mm) and quantification of infarct volume after MCAO from *Asic1a*^+/+^ and *Asic1a*^-/-^ mice applied with CGS19755 (1 mg/kg, i.p.), PcTX-1 (100 ng/kg, i.n.) and LK-2 (10 mg/kg and 30 mg/kg, i.p.). *n*=8, 12, 14, 6, 10 and 11 mice for each group. Data are mean±s.e.m.; one-way ANOVA with Tukey post hoc correction (**f**,**g**,**i**). *P* values are indicated.

To directly investigate the effects of LK-2 in a mouse stroke model *in vivo*, we first tested whether LK-2 could pass through BBB by using pharmacokinetics measurements. Injection of LK-2 at a dosage of 30 mg/kg (i.p.) led to a maximal concentration of 92660 μg/L in plasma and 236 μg/L in the brain, demonstrating that LK-2 could penetrate through the BBB (Extended Data Table.4) and reach a concentration in cerebrospinal fluid that is close to its IC_50_ for blocking glutamate-enhanced I_ASICs_ (Extended Data Table 4). In addition, LK-2 had a terminal elimination halflife (t_1/2_) of 2.297 hours, which theoretically constitutes a critical window of intervention for its neuroprotective effects after the onset of stroke (Extended Data Table 4 and Extended Data Fig. 11e). These results indicated that LK-2 is a potent small-molecule that displays favorable properties to provide neuroprotection against ischemic brain injury in *vivo*.

Using the mouse model of ischemic stroke induced by MCAO, we assessed the infarct volume with and without injection of LK-2 at the time of reperfusion (i.e. 30 min after artery occlusion). We observed a significant reduction in brain damage after administration of LK-2 at a dosage of 30 mg/kg compared to that of the saline group in *Asic1a*^+/+^ mice (Fig. 4h, i). Furthermore, the protective effect provided by LK-2 (30 mg/kg) against brain damage was comparable to that provided by CGS19755 (1 mg/kg, i.p.) (P=0.8288). Interestingly, *Asic1a*^-/-^ mice were resistant to brain injury by the MCAO paradigm and further administration of CGS19755 (1 mg/kg, i.p.) in these knockout mice added little to the extent of neuroprotection (Fig. 4i), reinforcing the indispensable role of glutamate-dependent potentiation of ASICs in mediating ischemic brain injury. Taken together, these results demonstrated that LK-2 functioned as a competitive antagonist for glutamate binding site on ASICs, providing neuroprotection by specifically abolishing glutamate-induced potentiation of I_ASICs_, independent of its action on NMDARs.

## Discussion

In this study, we demonstrated that glutamate and its structural analogs (e.g. agonists for NMDARs: NMDA, aspartate or competitive antagonist for NMDARs: L-AP5/APV) can function as allosteric positive modulators to potentiate currents mediated by ASICs. The association of glutamate with ASIC1a appears to play an overwhelming role in mediating NMDAR-independent neuronal injury through elevation of [Ca^2+^]_i_ and mitochondrial dysfunction in ischemic brain damage. Our *in silico* modeling and site-directed mutagenesis analyses identified an extracellular pocket of *h*ASIC1a with K380 being a critical amino acid residue for glutamate to directly bind to. We further provided a proof-of-principle that targeting the glutamate binding site on ASICs rather than NMDARs was successful in alleviating cell death *in vitro* and ischemic brain injury *in vivo*, with effective compounds such as LK-2 stemming from computer aided drug design based on the structure of CGS19755, another competitive antagonist of NMADRs (Extended Data Fig.12).

To date, strategies for developing neuroprotectants using NMDAR antagonists including CGS19755 (or *Selfotel*) appeared to be effective in animal models of stroke but have failed to provide protection in clinical trials or been terminated due to severe side effects including neuropsychiatric and tolerance issues^34,36–38^. NMDAR antagonists may not only block pro-survival functions of NMDARs during stroke recovery but also induce severe psychosis^39^, making them difficult targets for developing effective therapy. Consequently, enormous efforts have been made to unravel and target signaling pathways and protein substrates that are downstream to NMDARs to alleviate excitotoxicity in ischemic brain injury. Among previous studies, TRPM2, TRPM4 and ASIC1a channels have been shown to directly interact with NMDARs^5–7^. The mapping of their respective interaction domains provided means to disrupt NMDAR signaling without direct perturbations to the multifaceted roles of NMDARs in synaptic transmission and plasticity critical for cognitive function. All of this has converged towards the concept of NMDAR-dependent signaling underlying cell death. By contrast, we demonstrated that glutamate at clinically relevant concentrations under ischemic conditions potentiated I_ASICs_ and largely accounted for NMDAR-independent neurotoxicity in *vitro* and in *vivo*. Indeed, ASIC1a is responsible for glutamate-independent, acidosis-mediated ischemic brain injury at a low pH 6.5-6.0^4,40,41^. Interestingly, we found that potentiation of ASIC1a currents by glutamate at a mild pH range (e.g. 7.0-6.5) sensitizes neurons to such pH changes and boosts Ca^2+^ influx, membrane excitability and downstream signaling, driving cell death upon ischemic insult. The high sensitivity of ASIC1a to glutamate observed at pH range of around 7.0 can be fully accounted for by an increased affinity of ASICs for protons when glutamate occupies its co-agonist site on ASICs. Given that synaptic vesicles are loaded with extremely high concentration of glutamate and protons^42^, we favors the working model that the proton-ASIC1a-glutamate complex may be directly in play upon the onset of ischemic stroke, instead of being secondary to or downstream from the activation of NMDARs.

We demonstrated that glutamate directly binds to a novel cavity on the extracellular domain of ASIC1a, where both glutamate and protons can function as the first messengers. This is important because our findings highlight an unexplored signaling modality in the nervous system. Recent studies have shown that ASIC1a-containing channels can contribute to evoked EPSCs that were previously thought to be exclusively mediated by glutamate receptors in cortical neurons during synaptic transmission^43–45^. Therefore, our findings may have important ramifications for understanding the unappreciated roles of ASIC1a in synaptic transmission and plasticity underlying learning, memory and other behaviors^46–48^.

Of note, we also showed that a classical competitive NMDAR antagonist, CGS19755, abolishes glutamate-induced potentiation of I_ASICs_ without affecting basal function of ASICs. Its dual blockage of both ASICs and NMDARs likely resulted in the minimal infract volume as we and others have observed. To mitigate the potential psychotic issue of NMDAR antagonists while preserving the pro-survival role of NMDARs, we discovered a small molecule LK-2 based on the structure of CGS19755. Unlike classical ASICs blockers with poor selectivity or BBB permeability in drug delivery^4,49^, LK-2 at low micromolar concentrations preferentially decoupled the co-agonist effect of glutamate on ASICs without compromising basal ASIC and NMDAR gating, showing a promising effect of neuroprotection by selectively targeting the glutamate binding site on ASICs. Together, our findings build a new conceptual paradigm for mechanistic understanding of neurotoxicity in ischemic stroke and for strategizing the development of new stroke therapeutics that work independently from NMDARs.

## Methods

### Animals

Wildtype C57BL/6j mice were purchased from Jiesijie company (China) and housed in groups, in the animal facility at Shanghai Sixth People’s Hospital, Shanghai Jiaotong University. *Asic1a ^-/-^*(with congenic C57BL/6j background) mice are provided by Dr. Tianle Xu Lab. All mice were kept in standard cages (15 × 21 × 13.5 cm) on a 12:12 h light:dark cycle with ad libitum access to food, water, and nesting material. Animals were checked daily and their weight was monitored during the experiments. Animals were randomly allocated to treatment groups.

### Ethical approval

Throughout the study, all efforts were carried out to minimize animal suffering and reduce the number of animals. Experiments were conducted in conformity with the institutional principles for the care and use of animals, and experimental protocols that were approved by the Ethics Committee of Shanghai Sixth People’s Hospital.

### Electrophysiology from cultures and slices

#### Whole-cell recordings

Whole-cell patch clamp recordings were made from either neurons or CHO cells in a recording chamber (Corning) mounted on a fixed-stage upright microscope (Nikon). Patch electrodes (4-6 MΩ) were made from 1.5 mm borosilicate glass (World Precision Instruments). Whole-cell currents were recorded using an EPC-10 patch-clamp amplifier (HEKA). Data were acquired at 10-20 kHz and filtered at 1-3 kHz using a computer equipped with the Pulse 6.0 software (HEKA, Lambrecht). Cells were recorded at a holding potential of -60 mV unless otherwise described. For NMDAR current recordings, holding potential was -70 mV. A multi-barrel perfusion system (SF-77B, Warner Instruments) and pressure regulator system (ALA-VM8, Scientific Instrument) were used to achieve a rapid exchange of extracellular solutions. The perfusion protocol was set and controlled by the PatchMaster software. To avoid use-dependent desensitization or rundown, ASICs were repeatedly activated by acidic solution at least every 1 min. During each experiment, a voltage step of -10 mV from the holding potential was applied periodically to monitor the cell capacitance and access resistance. Dose-response curves were fitted to the Hill equation: *a=I/Imax=1/[1+10^n(b-EC^_50_)],* where *a* is the normalized amplitude of the I_ASICs_, *b* is the concentration of proton in external solution ([H^+^]), *EC_50_* is the [H^+^]-yielding half of the maximal peak current amplitude, and *n* is the Hill coefficient. The IC_50_ values for blocker dose-response curves were fitted using the following equation: *I/Imax=1/{1+[IC_50_/(blocker concentration)]^n^}*, where n is the Hill coefficient and *IC_50_* is the concentration of blocker producing 50% of the maximal block (*Imax*).

#### Single-channel recordings

Unitary currents were recorded using the outside-out configuration of the patch-clamp technique in ASIC1a transfected CHO cells. Channels were activated by rapidly moving squared-glass tubes delivering solutions of desired pH in front of the tip of the patch pipette. The delivery device achieves complete solution changes within 20 ms (SF-77B). When filled with solutions, pipettes had resistances of 6-10 MΩ. Single-channel currents were recorded with an EPC-10 patch-clamp amplifier (HEKA). The data were collected at 10 kHz and gain value 200 mV/pA, filtered at 1 kHz, and stored on a computer for analysis. Data were filtered off-line with a digital Gaussian filter to 1 kHz. All points amplitude histograms of ASIC unitary currents are fitted to the two-exponential equation:

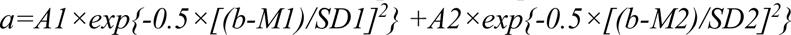

where *a* is the count of ASIC unitary currents, *b* is the amplitude of ASIC unitary currents. *A1* and *A2* are the heights of the center of the distribution, *M1* and *M2* are the amplitude of ASIC unitary currents at the center of the two distributions. *SD1* and *SD2* are measures of the widths of the distributions, in the same units as *b* (pA).

#### Solutions and chemicals

The artificial cerebrospinal fluid (aCSF) contained (in mM): 124 NaCl, 5 KCl, 1.2 KH_2_PO_4_, 1.3 MgCl_2_, 2.4 CaCl_2_, 24 NaHCO_3_, and 10 glucose, with pH 7.4 (300-330 mOsm). The extracellular fluid (ECF) for culture cells or neurons contained (in mM): 140 NaCl, 5 KCl, 1 CaCl_2_, 1 MgCl_2_, 10 glucose and 25 HEPES. For solution with pH≤6.8, MES was used instead of HEPES for stronger pH buffering. For NMDAR current recordings, MgCl_2_ was removed. Intracellular solution for voltage-clamp recordings contained (in mM): 120 K-gluconate, 2 MgCl_2_, 1 CaCl_2_, 10 HEPES, 0.5 EGTA, 4 Mg-ATP, with pH 7.3, and the osmolality was adjusted to 290-300 mOsm. For isolated neuron and NMDAR current recordings, K-gluconate was replaced by CsCl. All experiments were performed at room temperature (21-26°C). D-AP5, CGS19755, CNQX, Amiloride, PcTX-1 were purchased from Alomone Lab, L-AP5 and CGP39551 were purchased from Tocris, and other chemicals were purchased from Sigma-Aldrich. It should be noted that all ECF pH in this study added with any chemicals were adjusted to indicated values (pH changes less than 0.02 after adding chemicals).

#### CHO cell culture and transfection

Chinese hamster ovary (CHO) K1 cells were cultured in F12 medium with 10% FBS (Gibco). Penicillin/streptomycin (Invitrogen) were added to the medium for preventing bacteria contamination at a final 1% concentration. For NMDAR expression, 500 μM D-AP5 was added in the cultured medium to avoid glutamate-induced toxicity via activating NMDAR^1^. 0.25% Trypsin-EDTA (Gibco) was used for cell passage. Cells were incubated at 37°C in a humidified CO_2_ incubator.

CHO cells were transfected with plasmids described in detail previously^2^. Briefly, at a cell density of 50%-70%, a total 3.0 μg cDNA mixed with Lipofectamine 3000 transfection kit (Invitrogen) was added to 35 mm dishes. For co-expression of ASIC1a plus 2a or 2b, equal amounts of cDNA for both subunits were used. For NMDAR expression, plasmids of human GluN1, GluNR2A or GluNR2B and PSD-95 at a ratio of 1:4:0.5 were used. cDNA of green fluorescent protein (GFP) was linked at the N-terminus of those proteins. All experiments were performed 24-48 hours after transfection, all dishes were washed three times using ECF before experiments, and GFP-positive CHO cells were viewed under a fluorescent microscope for patch-clamp recordings. Human and mouse ASIC subunits were used for constructing all wildtype and mutants in this study. Site-directed mutagenesis was performed on the WT plasmid using Takara PrimeSTAR^TM^ HS DNA polymerase.

#### Primary cortical neuron culture

Primary cortical neurons were prepared and maintained as previously described^3^. Briefly, cerebral cortices from 24 hours postnatal *Asic1a^+/+^* or *Asic1a^-/-^*mice were dissected in DMEM high glucose solution and dissociated by 0.05% trypsin for 15 min.

Cells were plated (∼2×10^5^ cells/35 mm dish for electrophysiology and immunocytochemistry) on poly-D-lysine coated cover glasses or dishes. Cultures were maintained in Neurobasal-A medium (Gibco) containing 2% B27 (Gibco) and 1% Glutamax supplements (Gibco) at 37°C in a 5% CO_2_ humidified atmosphere.

#### Acute isolation of cortical neurons

Acute dissociation of mouse cortical neurons was performed as described previously^4^. Briefly, mice from postnatal day 15 to 18 were anaesthetized with isoflurane. Cortical tissues were dissected and incubated in oxygenated ice-cold aCSF (with half concentration of CaCl_2_). Transverse cortical slices (500 µm) were cut with a microtome (Leica VT1200) followed by incubation in aCSF containing 3.5 mg ml^-1^ papain (Sigma-Aldrich) at 37°C for 30 min. Slices were then washed three times and incubated in enzyme-free ECF solution for at least 15 min before mechanical dissociation. For dissociation, slices were triturated using a series of fire-polished Pasteur pipettes with decreasing tip diameters. Recording began 15 min after the mechanical dissociation. Only the neurons that retained their pyramidal shape and dendrites were used for recordings.

#### Acute brain slices preparation

Brain slices were prepared from 6 to 8-week-old C57BL/6j *Asic1a^+/+^*or *Asic1a^-/-^* mice that were anesthetized with pentobarbital (55 mg/kg, i.p.) and decapitated. The brain was quickly removed and submerged into oxygenated ice-cold aCSF. The transverse brainstem slices were sectioned at 250 μm (for bioassay) with a microtome (VT-1200S, Leica). The slices were then incubated in aCSF at temperature 37 °C for at least 30 min. All solutions were saturated with 95% O_2_-5% CO_2_.

### Ca^2+^ imaging

Primary cortical neurons were incubated in a confocal microscopy (Zeiss 710) specialized dish (35×35 mm, Cellvis) with ECF (saturated with 95% O_2_) and 1 μM fluo-3 AM (Beyotime) for 20 min at room temperature, followed by three times wash and additional incubation in normal ECF for 15 min. The dish was then transferred onto the stage of confocal microscopy. Neurons were illuminated using a xenon lamp and observed with a 40×UV fluor objective lens. The shutter and filter wheel were set to allow for 488 nm excitation wavelength. Images were analyzed by every 5 second in circumscribed regions of cells in the field of view. Digital images were acquired, stored and analyzed by the ZEN software (Zeiss). For quantification, intracellular calcium levels were plotted as ΔF/F_0_ ratios over time, where F_0_ is the initial fluorescence intensity of each cell.

### Imaging of mitochondrial membrane potential

Mitochondrial membrane potential (Ψm) was measured using a JC-1 kit (Beyotime). Primary cortical neurons were loaded with JC-1 working solution in for 20 min, then washed and incubated in JC-1 washing buffer for 5 min before recording. JC-1 was imaged with 488 nm and 546 nm excitation wavelengths using 40× objective. For quantification, Ψm was measured as the fluorescence intensity ratio of red and green (R/G), then the ratio was normalized to the initial fluorescence intensity ratio of each cell.

### Calcein-PI cell death assay

Acute isolated brain slices (250 μm) were incubated for 1 hour at room temperature with different treatment and followed by exposure to 1 μM calcein-am and 2 μM PI (Propidium-Iodide) (Solarbio) for 20 min (solutions were saturated with 95% O_2_-5% CO_2_ in the process). Then brain slices were washed with ECF for three times. After fixation in 4% paraformaldehyde (Solarbio) for 40 min (covered by a light-resistant container), slices were observed shortly after staining by confocal microscopy. The live and apoptotic nuclei was determined by 488 nm and 546 nm excitation wavelengths using microscopic examination at 5× or 40 × magnification. The number of staining nuclei was counted by using the Image-J software. The percentage of cell death was calculated as: *cd%=[nred/(nred+ngreen)]*100,* where *cd%* is the percentage of cell death, *nred* and *ngreen* are the number of PI (dead cells) and calcein (live cells) staining cells, respectively.

### Cell injury assay with LDH measurement

Brain slices were washed three times with ECF and randomly divided into treatment groups and incubated in different solutions at room temperature for 1 hour. LDH release was measured in ECF using LDH assay kit (Beyotime). Following incubation, culture medium (120 μl) was transferred to 96-well plates and mixed with 60 μl reaction solution provided by the kit. Optical density was measured at 490 nm and 620 nm 30 min later, using a microplate photometer (BioTek). The maximal releasable LDH in each group was obtained by incubation with 1% Triton-100.

### Middle cerebral artery occlusion (MCAO)

Transient focal ischemia was induced by suture occlusion of the middle cerebral artery (MCAO) in *Asic1a^+/+^* and *Asic1a^-/-^* mice. Animals were anesthetized using an animal mini anesthesia machine (RWD). During anesthesia induction, animals were put in a chamber (15×25×30 cm) with 2% isoflurane; animals were put on a mask with 1% isoflurane during MCAO operation. Adequate ischemia was confirmed by continuous laser Doppler flowmetry (moor FLPI-2). Animals that did not have a significant reduction of blood flow less than 50% baseline values during MCAO were excluded. Rectal and temporalis muscle temperature was maintained at 33±0.5°C with a thermostatically controlled heating pad (Extended Data Table 5). Intraperitoneal (i.p.) injection was performed immediately after removing the suture occlusion. Animals were killed with isoflurane overdose 24 h after ischemia.

Brains were removed and dissected coronally at 1 mm intervals, and stained with the vital dye 2,3,5-triphenyltetrazolium hydrochloride (TTC) and the normal area will be stained with TTC. Infarct volume (%) was calculated by summing infarction areas of all sections and multiplying by slice thickness, then dividing the whole volume of the brain. Manipulations and analyses were performed by individuals blinded to treatment groups. Depending on the experimental design, 30 min MCAO was performed for moderate ischemic model.

### Laser speckle imaging

Mice were anaesthetized by 1% isoflurane and their heads were restrained in a stereotaxic cylinder frame to minimize breathing motion. The scalp and the skull fascia were gently incised down the midline and peeled to the side. Saline was titrated onto the skull to maintain moist. Laser speckle images were recorded with a CMOS camera before MCAO, 15 min after occlusion and 15 min after reperfusion. For each animal, three sets of raw speckle images were acquired in <15 s (250 frames in each set; image width, 752 pixels; image height, 580 pixels; exposure time, 20 ms). A speckle contrast image was calculated from each raw speckle image using a sliding grid of 2.5 mm × 2.5 mm. A mean speckle contrast image was calculated for each set and used to calculate the relative cerebral blood flow (rCBF). The rCBF in the ipsilateral (ischemic) hemisphere was normalized by the mean rCBF in the contralateral (non-ischemic) hemisphere. Speckle images were obtained and processed by the software mFLPI2MeasV2.0, rCBF data from all pooled hemispheres were obtained by the software moorFLPIReviewV50. All analyses were randomized.

### Molecular docking and Prime-MM/GBSA binding free energy calculation

Structures of cASIC1a (PDB ID: 5WKU, resting state) and NMDAR (PDB ID:5IOU) were obtained from Protein Data Bank. Structure of chemicals was obtained from PubChem compound or ChemBioDraw Ultra 14 software. Initial docking studies involved preparation of the cASIC1a and glutamate using the High-Ambiguity Driven protein-protein DOCKing (HADDOCK) software (https://www.bonvinlab.org/software/haddock2.4/). Initial grids included the entire extracellular surface of the protein. After docking calculations, the highest scoring conformations were analyzed, the results revealed glutamate-binding in six extracellular residues (Extended Data Table 1). Following identification of a general binding site, additional docking poses were generated for the most potent binding sites using the Schrödinger Maestro software suite (Schrödinger, 2020-3). Prior to docking, protein was processed using the Protein Preparation Wizard for adding missing residues, optimizing side-chains, removing waters, optimizing H-bond and energy (OPLS3e force field). Ligand was optimized using OPLS3e force field in LigPrep module. Protein and ligand protonation states at pH 7.0±0.2 were sampled using Epik.

Ligand was docked to a picked residue (e.g. K379) in a grid box with dimensions of 25 × 25 × 25 Å^3^. Extra-precision docking (Glide XP) was performed with flexible ligand sampling, and post-docking minimization was performed to generate a maximum of 10 poses per ligand within the Glide program, the docking conformation with a highest docking score was analyzed. The binding free energies of all different poses from XP docking outputs were carried out using Prime-MM/GBSA module^5^. The binding energy was calculated by the software according to the following equation:

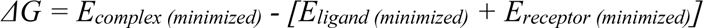

### Molecular dynamics (MD) simulation

For MD simulation, the best pose of each ligand-protein complex was selected from docking results and the ligand-bound protein systems were built in 150 mM NaCl aqueous solution. To investigate the stability of the docked ligand-protein poses, 50-ns simulations were performed. After 25000 steps of minimization, the systems were equilibrated using isothermal-isobaric (NPT) ensembles at a constant temperature of 303.15 K, followed by 50-ns production run. All simulations were using the GROMACS 2020.3, CHARMM36m force fields for the protein, and GRO-MOS 54A7 force field for ligand^6^, and the SPCE model for water. The simulation trajectories were analyzed for structural stability using root-mean-square deviation (RMSD) and root-meansquare fluctuation (RMSF) calculations. MM-PBSA calculations on MD simulation trajectories were performed with a modified gmx_mmpbsa bash script (available at https://github.com/Jerk-win/gmxtools/tree/master/gmx_mmpbsa) using solvent-accessible surface area (SASA) as the model for non-polar solvation energy.

### CGS19755 binding pocket analysis and drug design

The best pose of CGS19755-cASIC1a complex was selected from molecular docking for binding pocket analysis by Fpocket 2.0 software. A dpocket program was performed to produce pocket parameter using default settings. We used a scaffold replacement method based on CGS19755 structure to screen the ZINC20 database (fragment, lead-like, and drug-like molecules) by the Molecular Operating Environment (MOE) software (2015.10). The three-dimensional (3D) conformations of the remaining about 222 compounds were generated by the ligPrep module of Maestro (Schrödinger) with the OPLS3e force field. Possible ionization states of each compound were generated in the pH range of 7.0±0.2 using Ionizer. Possible tautomer forms were also generated for each ligand. Compounds were screened using the high throughput virtual screening module followed by the extra precision docking module in Glide. The Glide docking score was used to rank the results list. Finally, 6 hits were manually selected for electrophysiological assay.

### Pharmacokinetics study

Pharmacokinetics of LK-2 was analyzed in male C57BL/6j mice (n=22). Plasma and brain concentrations were determined using LC-MS/MS methods after a single intraperitoneal injection dose (i.p. 30 mg/kg) of compound as a clear solution in 0.9% saline at a concentration of 1 mg/ml. Blood samples were collected into EDTA-coated test tube at the time point of 0.083 h, 0.25 h, 0.5 h, 1.0 h, 2.0 h and 4.0 h, and then centrifuged at 2,000 g for 15 minutes to generate plasma samples. Brain samples were collected after intra-ventricle perfusion with normal saline and prepared by homogenizing tissue with 5 volumes (w:v) of 0.9% NaCl. LC-MS/MS methods to quantify LK-2 in plasma and brain samples were developed with the instrument of LC-MS/MS-T_API4000. General sample processing procedure performed as following: 1) An aliquot of 30 µl plasma sample, calibration standard, quality control, single blank, and double blank sample was added to the 1.5 ml tube respectively; An aliquot of 40 µl brain homogenate, calibration standard, quality control, single blank, and double blank sample was added to the 96-well plate respectively; 2) Each sample (except the double blank) was quenched with 150 µl (for plasma samples) or 200 µl (for brain homogenates) of IS 1 [6 in 1 internal standard in MeOH (Labetalol & tolbutamide & Verapamil & dexamethasone & glyburide & celecoxib 100 ng/ml for each) with 40 mM DBAA)] respectively (double blank sample was quenched in MeOH with 40mM DBAA), and then the mixture was vortex-mixed well (at least 15 s) and centrifuged for 15 min at 12000 g (for plasma samples) or 3220 g (for brain homogenates), 4 °C; 3) An aliquot of 65 µl supernatant was transferred to the 96-well plate and centrifuged for 5 min at 3220 g, 4 °C. All the processes were done on the wet ice. All the supernatants were directly injected for LC-MS/MS analysis. The column used was an ACQUITY UPLC BEH C18 2.1 × 100 mm, 1.7 μm column. Column temperature was 40 °C. Flow rate was 0.4 ml/min. The mobile phase consisted of A: 0.001% NH_3_·H_2_O with 0.18 mM DBAA in water, and B: 10 mM DMHA and 3 mM NH_4_OAc in ACN/Water (v:v, 50:50). Standard curves were prepared by spiking compounds into control plasma and brain and these were used to determine drug concentrations. Pharmacokinetic parameters were calculated via non-compartmental analysis (NCA) using DAS version 2.0 with mean concentration at each time point.

## Statistics

All data were reported as mean ± S.E.M. Two-tailed paired and unpaired Student’s *t-*test were used where appropriate to examine the statistical significance of the difference between groups of data. Comparisons among multiple groups were analyzed by One-way and two-way ANOVA followed by Tukey or Dunnett multiple comparison tests for *post hoc* analysis. Statistical software Prism 8 was used to analyze all data.

## Acknowledgements

This work was supported by National Key R&D Program of China (2021ZD0201900) to S.K.Y.; Shanghai Municipal Education Commission-Gaofeng Clinical Medicine Grant Support (20152233), International Cooperation and Exchange of the National Natural Science Foundation of China (82020108008) and Shanghai Jiao Tong University School of Medicine Multicenter Clinical Research Program (DLY201823) to H.B.S.; the National Natural Science Foundation of China (82101234), China Postdoctoral Science Foundation (2020M681335) and Shanghai Sixth people’s hospital (ynqn202108) to K.L.; and Canadian Institutes of Health Research Project Grants (PJT-156034 and PJT-156439), Natural Science and Engineering Research Council Discovery Grant (RGPIN-2017-06665) and Tier 1 Canada Research Chair Program to L.Y.W. We thank Feng Tang and Le Liu at *Simcere Pharmaceutical* for synthesis of all candidate compounds and Dr. Adam Fekete for critical comments on earlier versions of the manuscript.

## Author contributions

L.Y.W., S.K.Y. and H.B.S. conceived and designed this study. K.L. performed electrophysiological recordings and computational simulations in collaboration with I.P. and J.F.K. Z.Q.L. and

L.N.G. performed histology and imaging. H.W.L and M.X.L. performed animal modelling and histology. J.F.L. and X.Q. performed cell culture and transfection. T.L.X. provided ASIC1a mutant constructs and intellectual inputs. K.L. and L.Y.W. wrote the manuscript.

---

**Extended Data Figure 1.**
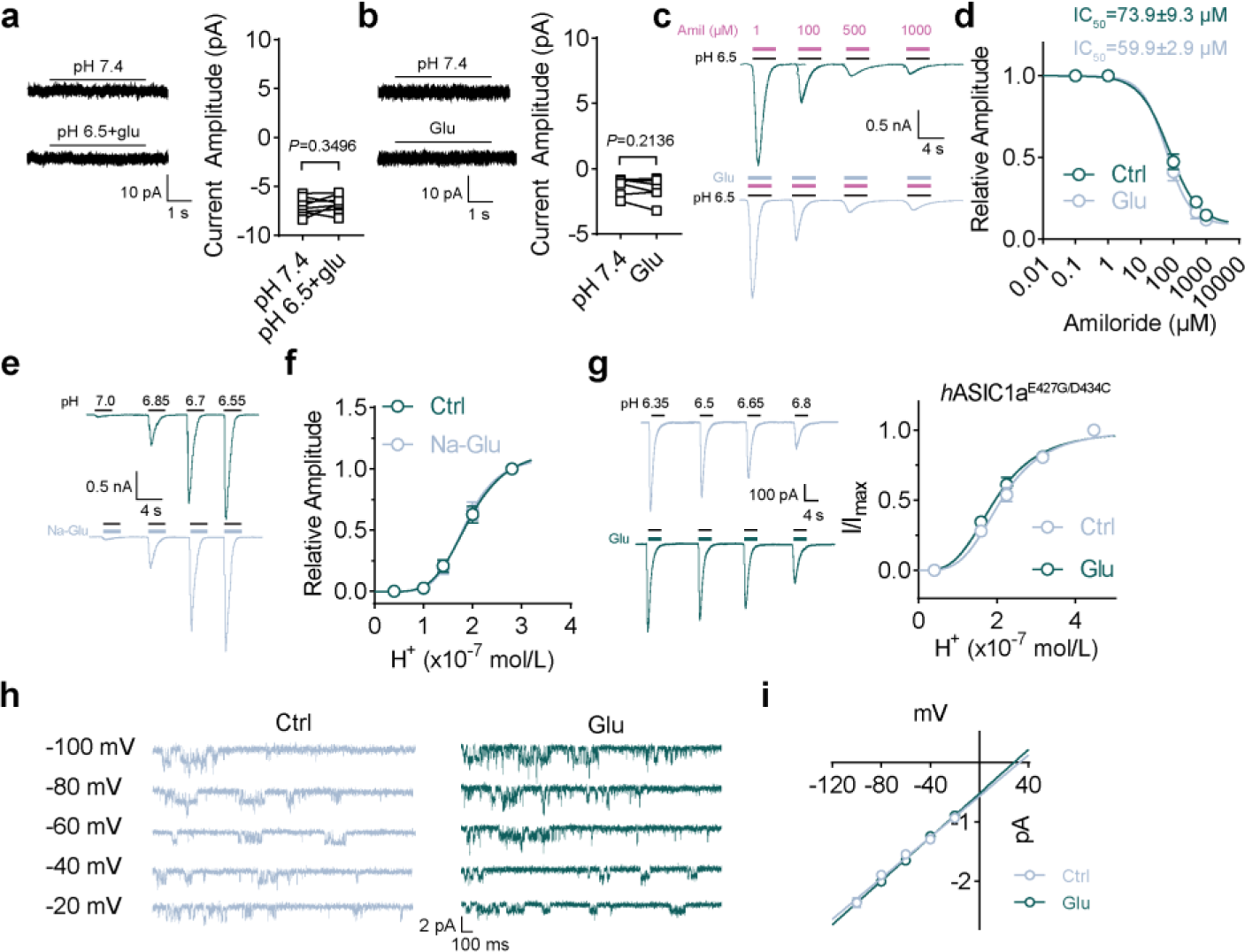
Glutamate binding to ASIC1a potentiates its current without altering its pharmacological properties and single-channel conductance. **a**, Glutamate (500 μM) did not activate currents in blank CHO cells even at pH 6.5. *n*=8 cells. **b**, Glutamate (500 μM) did not activate currents in ASIC1a transfected CHO cells at pH 7.4. *n*=6 cells. **c**,**d**, Inhibition of I_ASICs_ by amiloride at a dose-response manner was not affected by 1 mM glutamate. *n*=6 cells. **e**,**f**, Glutamic acid monosodium did not enhance I_ASICs_ by stepwise changes in solution with pH value from 7.4 to 7.0, 6.85, 6.7 and 6.55. *n*=5 cells. **g**, Representative traces and dose-response curves showing glutamate potentiated I_ASICs_ mediated by mutant *h*ASIC1a^E427G/D434C^ devoid of the Ca^2+^ binding sites. *n*=7 cells. **h**,**i**, Representative traces and current-voltage relationship showing ASIC1a unitary currents recorded at holding potential from - 100 mV to -20 mV (20 mV increment) in the presence and absence of 500 μM glutamate at pH 7.0. *n*=7 cells. Data are mean±s.e.m.; two-tailed paired Student’s *t*-test (**a**,**b**); *P* values are indicated.

**Extended Data Figure 2.**
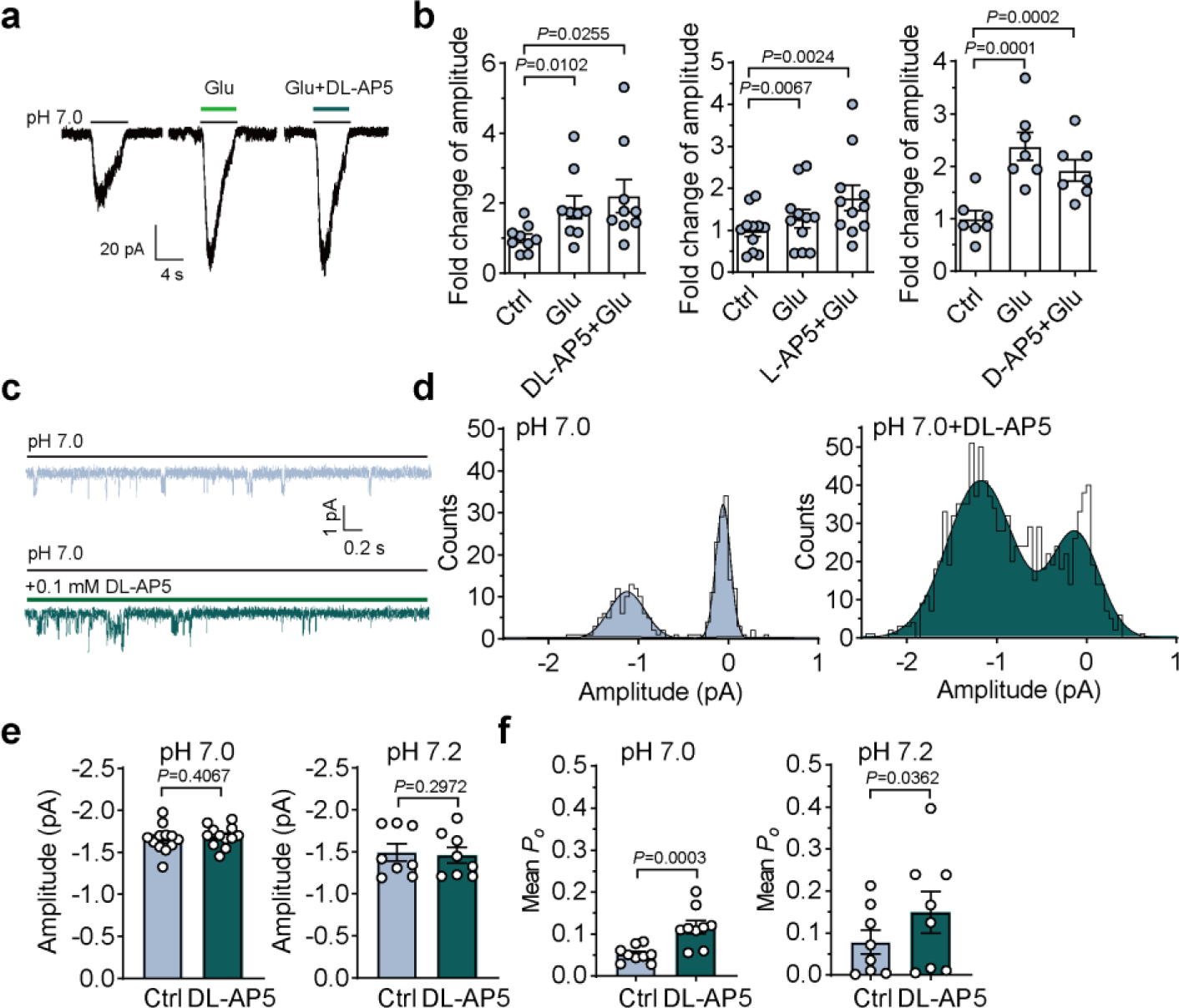
NMDAR antagonist AP5 does not block glutamate-dependent potentiation of ASIC1a currents and directly enhances open probability of ASIC1a single channel. **a**, Typical traces showing the effects of glutamate and DL-AP5 on I_ASICs_. **b**, Summary data showing DL-, L- and D-isomers of AP5 (200 μM) did not block glutamate-enhanced I_ASICs_. *n*=9, 11 and 7 cells. **c**, Outside-out patch recordings of ASIC unitary currents in an ASIC1a transfected CHO cell in the presence and absence of 100 μM DL-AP5 at pH 7.0. Currents were recorded at -60 mV. **d**, All points amplitude histogram of of ASIC unitary currents was constructed from (**c**), curves were fitted by two Gaussian components. Bin=0.05 pA. **e**,**f**, Quantification of amplitude and mean open probability (*P*_o_) of ASIC1a unitary currents evoked in the presence and absence of 500 μM glutamate at pH 7.0 and 7.2. *n*=12 and 8 cells for amplitude; *n*=9 and 8 cells for *P*_o_. Data are mean±s.e.m.; one-way ANOVA with Dunnett post hoc correction for multiple comparisons (**b**); two-tailed paired Student’s *t*-test (**e**,**f**); *P* values are indicated.

**Extended Data Figure 3.**
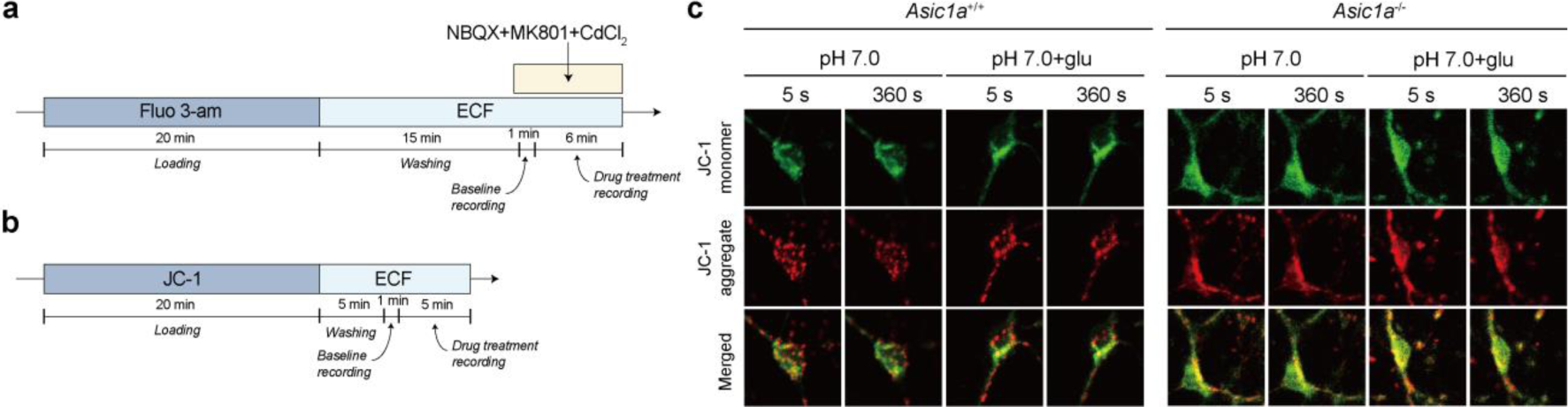
Ca^2+^ imaging and mitochondrial potential assay in cultured cortical neurons from *Asic1a*^+/+^ and *Asic1a*^-/-^ mice. **a**, Schematic of the experimental protocol of Ca^2+^ imaging. Cultured cortical neurons were loaded with Fluo 3-AM for 20 min followed by a 15 min washing step in extracellular fluid (ECF). After 1 min baseline recording, neurons were imaged for 6 min following pH 7.0 solution with or without glutamate in the presence of 10 μM NBQX, 1 μM MK801 and 100 μM CdCl_2_ to block AMPA receptors, NMDA receptors and voltage-gated calcium channels, respectively. **b**, Schematic flow of the experimental protocol of mitochondrial potential assay. Cultured cortical neurons were loaded with JC-1 for 20 min followed by a 5 min washout. After 1 min baseline imaging, neurons were recording for 5 min following pH 7.0 solution with or without glutamate. **c**, Representative images showing the mitochondrial potential changes with different treatment from wildtype and *Asic1a*^-/-^ mice. With mitochondrial potential (absolute value) decreasing, fluorescence intensity of JC-1 monomer (green) enhanced, while JC-1 aggregate (red) faded. First frame at 5 second and last frame at 360 second are shown here for comparisons.

**Extended Data Figure 4.**
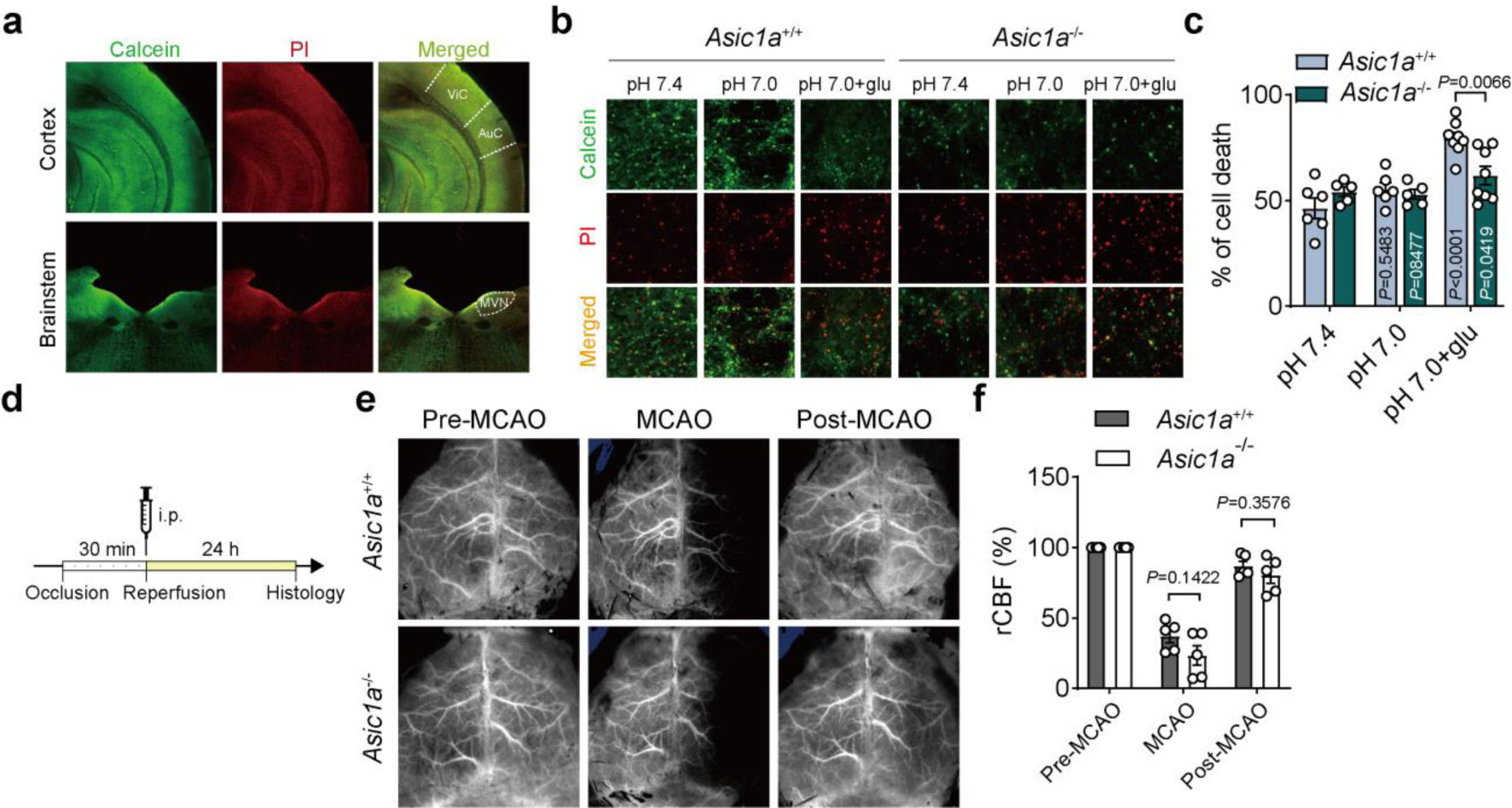
Calcein-PI staining of brain slices and Doppler laser speckle imaging for MCAO. **a**, Representative images showing coronal view of cortex and brainstem at low magnification under control condition (pH 7.4). ViC, visual cortex; AuC, auditory cortex; MVN, media vestibular nucleus. **b**,**c**, Representative images and summary data showing percentage of cell death by calcein-PI staining of brainstem (MVN area) slices from *Asic1a*^+/+^ (*n*=6, 6 and 8 slices for each group) and *Asic1a*^-/-^ (*n*=5, 5 and 8 slices for each group) mice under different conditions. Calcein (green), live cells; PI (red), dead cells. **d**, Schematic illustration of the timeline of MCAO treatment. **e**, Laser speckle imaging at 5 min pre-MCAO, 15 min post-occlusion (MCAO) and 15 min post-reperfusion (post-MCAO) in *Asic1a*^+/+^ and *Asic1a*^-/-^ mice. **f**, Summary data showing the relative cerebral blood flow (rCBF) changes during MCAO in *Asic1a*^+/+^ and *Asic1a*^-/-^ mice. *n*=5 mice for each group. Data are mean±s.e.m.; two-way ANOVA with Tukey post hoc correction for multiple comparisons (**c**,**f**); *P* values are indicated.

**Extended Data Figure 5.**
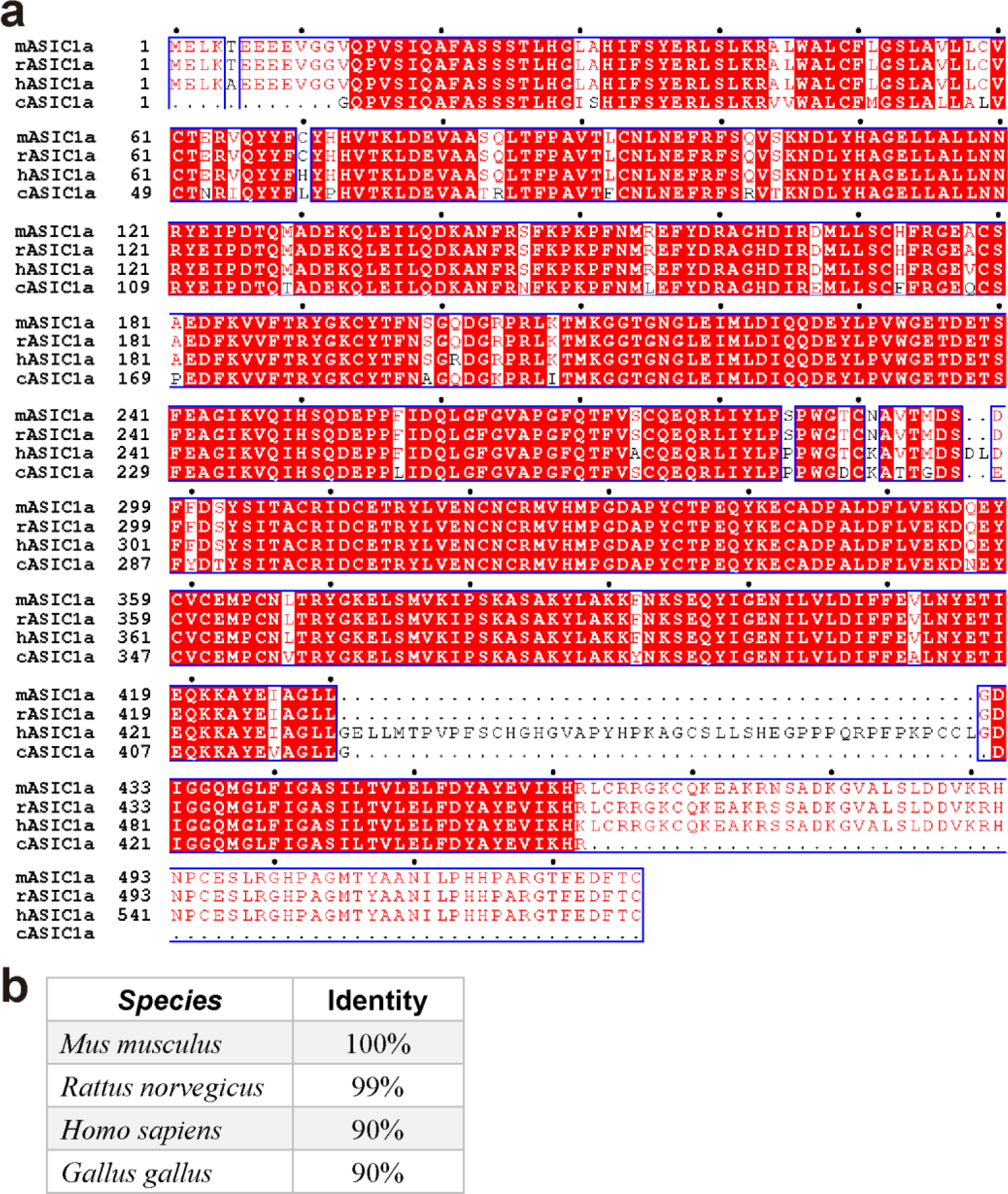
Alignment of ASIC1a among different species shows highly conserved sequences in *Aves* and *Mammalia*. **a**, Sequence alignment of the *m*ASIC1a (mouse), *r*ASIC1a (rat), *h*ASIC1a (human) and *c*ASIC1a (chicken). Homologous regions are colored red background. **b**, Table showing percentage of ASIC1a amino acid sequence identity among different species.

**Extended Data Figure 6.**
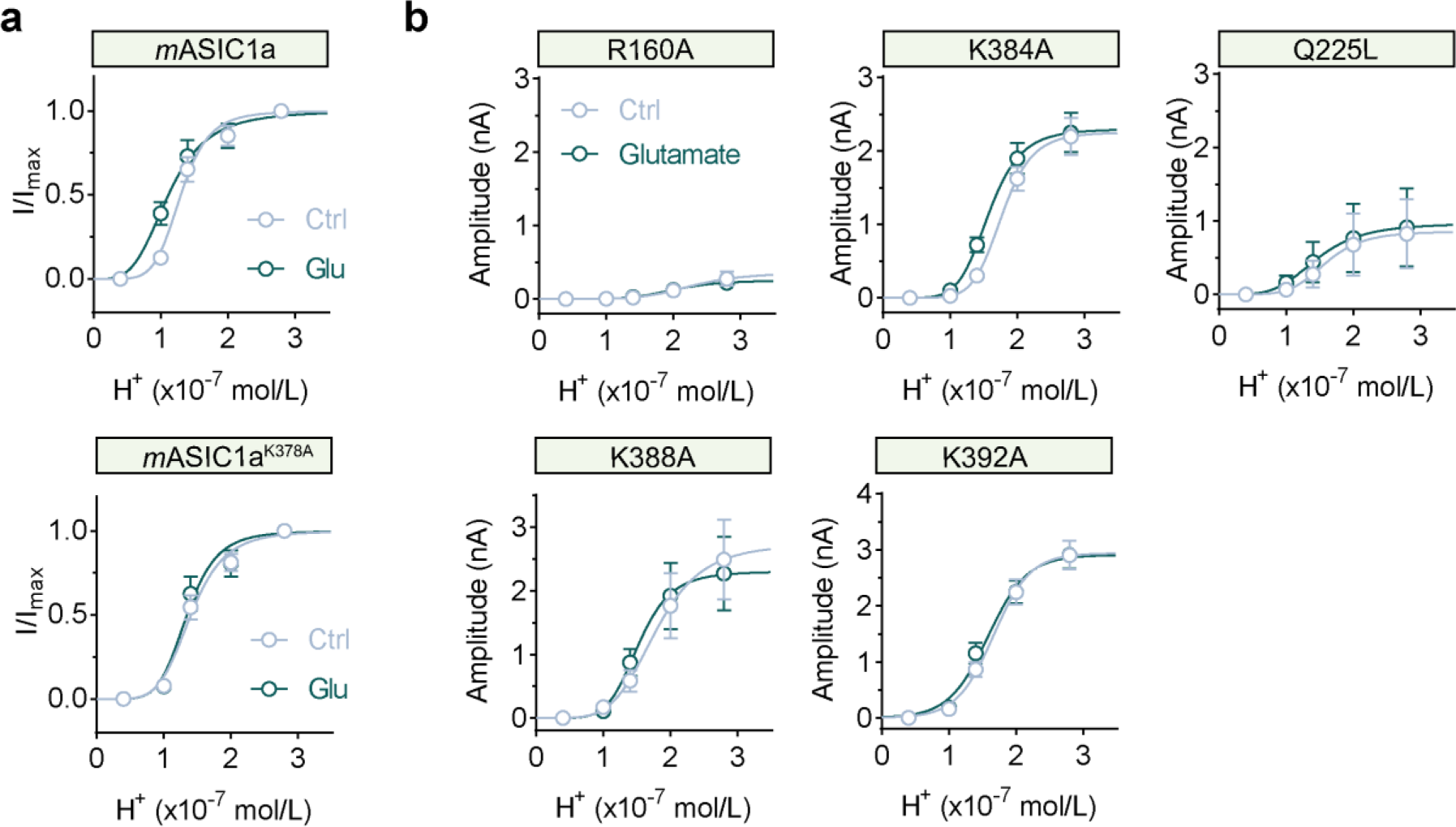
Screening glutamate-binding sites by site-directed mutagenesis. **a**, Dose-response curves showing the effects of 500 μM glutamate on *m*ASIC1a (*n*=5 cells) and *m*ASIC1a^K378A^ (*n*=6 cells) currents in CHO cells. **b**, Dose-response curves showing the effects of 500 μM glutamate on I_ASICs_ in R160A (*n*=5 cells), K384A (*n*=7 cells), Q225L (*n*=3 cells), K388A (*n*=9 cells) and K392A (*n*=15 cells) mutants of *h*ASIC1a transfected CHO cells. Glutamate enhances I_ASICs_ in K384A, Q225L and K388A mutants transfected CHO cells but not in R160A and K392A mutants. However, I_ASICs_ in R160A mutant significantly decreased in amplitude when compared to that by *m*ASIC1a, making it unlikely to be a glutamate-binding site. Data are mean±s.e.m.

**Extended Data Figure 7.**
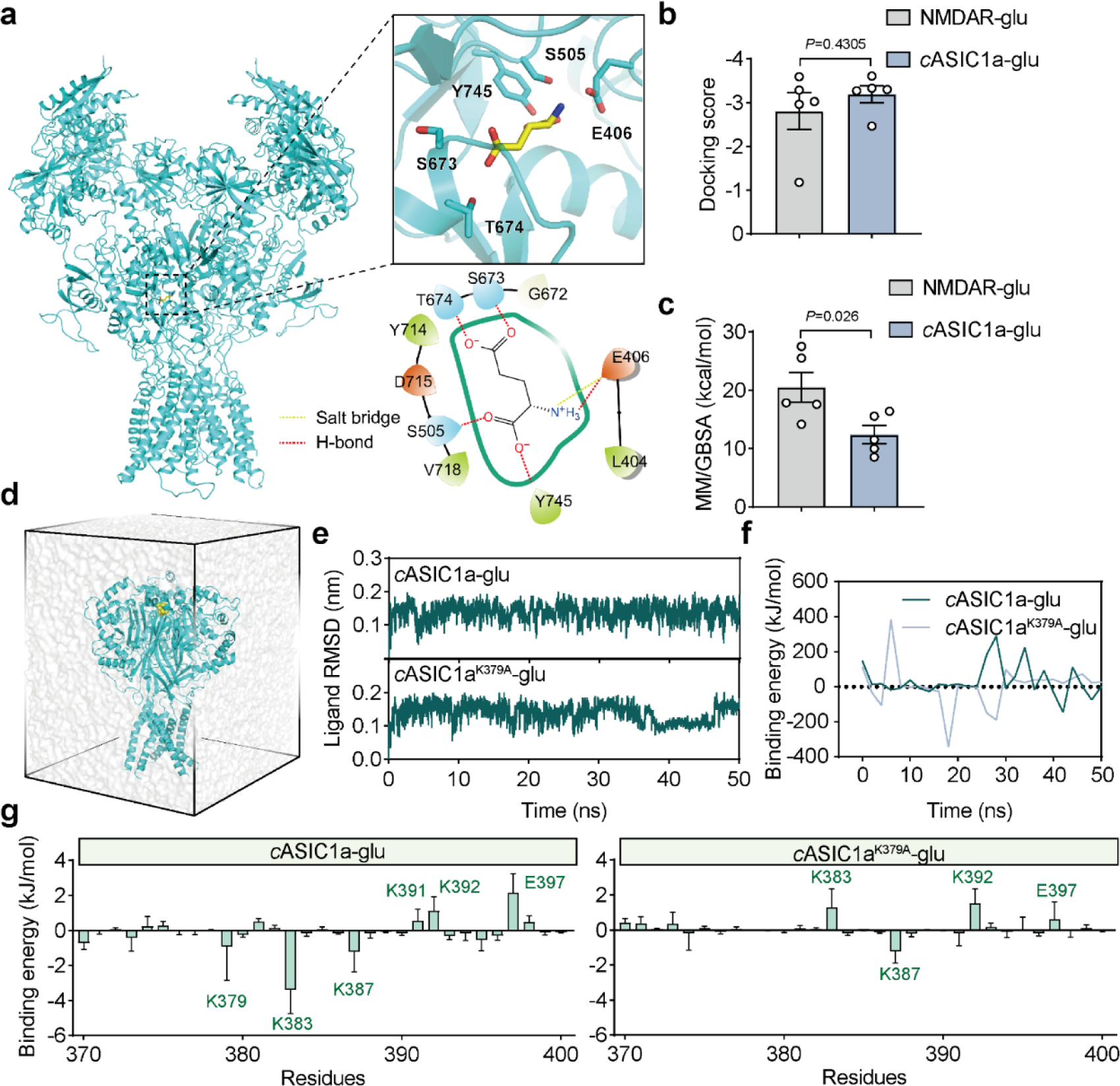
In *silico* analyses for glutamate-bounded NMDAR and *c*ASIC1a identify key residues forming the glutamate binding pocket on ASIC1a. **a**, Overall structure of glutamate-NMDAR complex (PDB: 5IOU) and a close-up view of glutamate-binding pocket. The interactions between glutamate and surround residues are shown as yellow (salt bridge) and red (hydrogen-bond) dash lines. **b**,**c**, Summary data by computational calculation showing docking scores and MM/GBSA of glutamate binding to NMDAR and *c*ASIC1a. *n*=5 and 5 poses, respectively. **d**, A snapshot of the simulation box of the glutamate-*c*ASIC1a complex in 150 mM NaCl solution. Glutamate is shown as yellow spheres, *c*ASIC1a is shown as cyan cartoon, water is shown as transparent surface. For clarity, ions are omitted. **e**, Structural stability of ligand in wildtype and mutant conformations was measured as the RMSD (unit: nm) over a 50-ns time course. **f**, Binding energy for glutamate-*c*ASIC1a complex was calculated by the MM-PBSA method. **g**, Binding energy of glutamate-*c*ASIC1a complex for amino acid residues from 370 to 400 over a 50-ns time course. Several residues with high binding energy were labeled. Data are mean±s.e.m.; two-tailed unpaired *t*-test (**b**,**c**).*P* values are indicated.

**Extended Data Figure 8.**
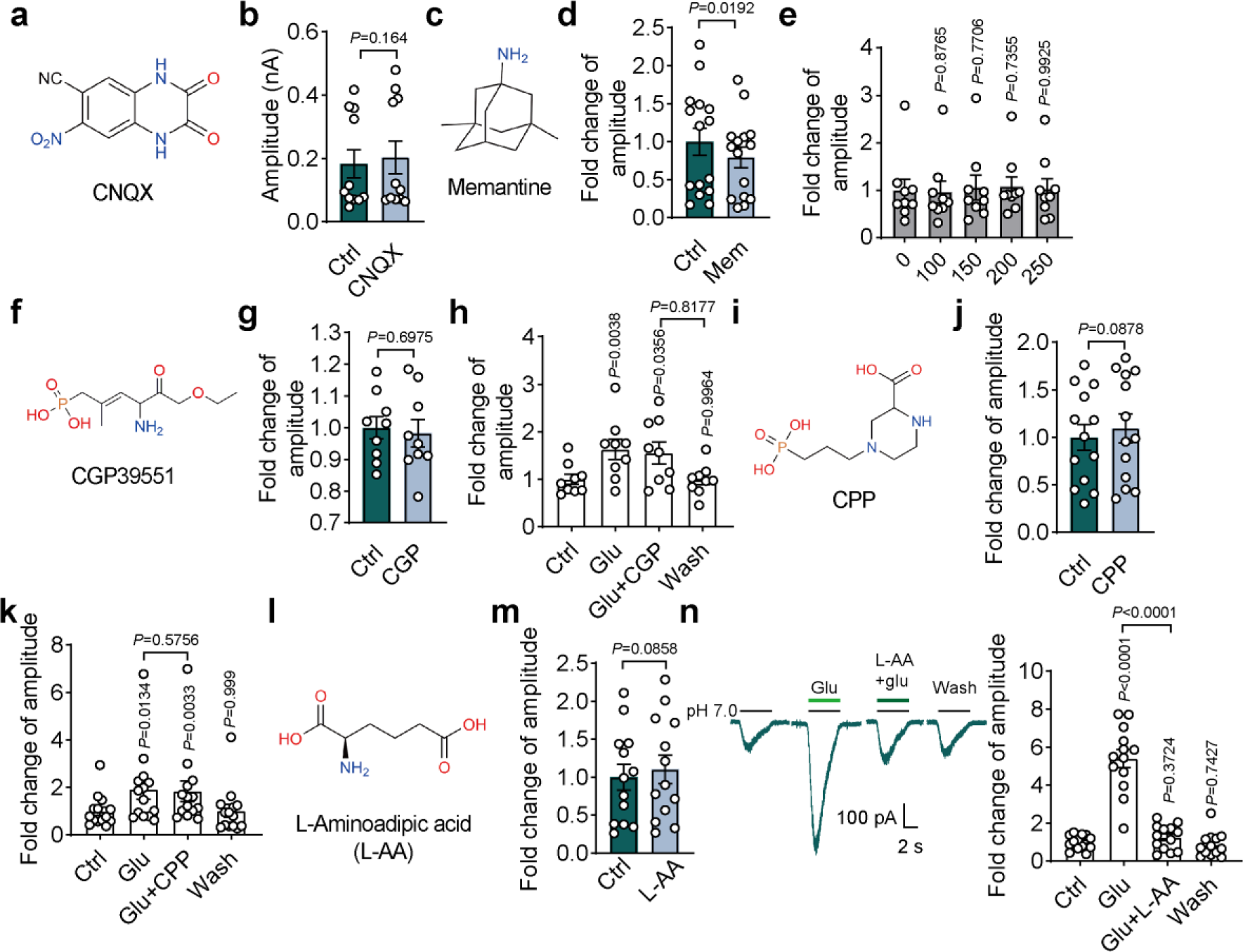
Pharmacological effects of glutamate receptor antagonists and agonist on I_ASICs_ in CHO cells. **a,c,f,i,l,** Chemical structure of CNQX, Memantine, CGP39551, CPP and L-Aminoadipic acid (L-AA). **b**, Summary data showing 10 μM CNQX, an AMPA receptor antagonist, had no effect on I_ASICs_. *n*=11 cells. **d**,**e**, Memantine, an open channel blocker of NMDA receptors, had no effect on I_ASICs_. *n*=15 cells (**d**) and 9 cells (**e**). **g**,**h**,**j**,**k**, 100 μM CGP39551 and CPP, competitive antagonists of NMDA receptors, had no effect on I_ASICs_ and cannot block glutamate-induced potentiation of I_ASICs_. *n*=9 cells (**g**), 9 cells (**h**), 13 cells (**j**) and 14 cells (**k**). **m**,**n**, 100 μM L-AA, an agonist for NMDAR and metabotropic glutamate receptors (mGluRs), had no effect on I_ASICs_, however, can eliminate glutamate-induced potentiation of I_ASICs_. *n*=13 cells (**m**) and 13 cells (**n**). Data are mean±s.e.m.; two-tailed paired *t*-test (**b**,**d**,**g**,**j**,**m**); one-way ANOVA with Tukey post hoc correction (**e**,**h**,**k**,**n**). *P* values are indicated.

**Extended Data Figure 9.**
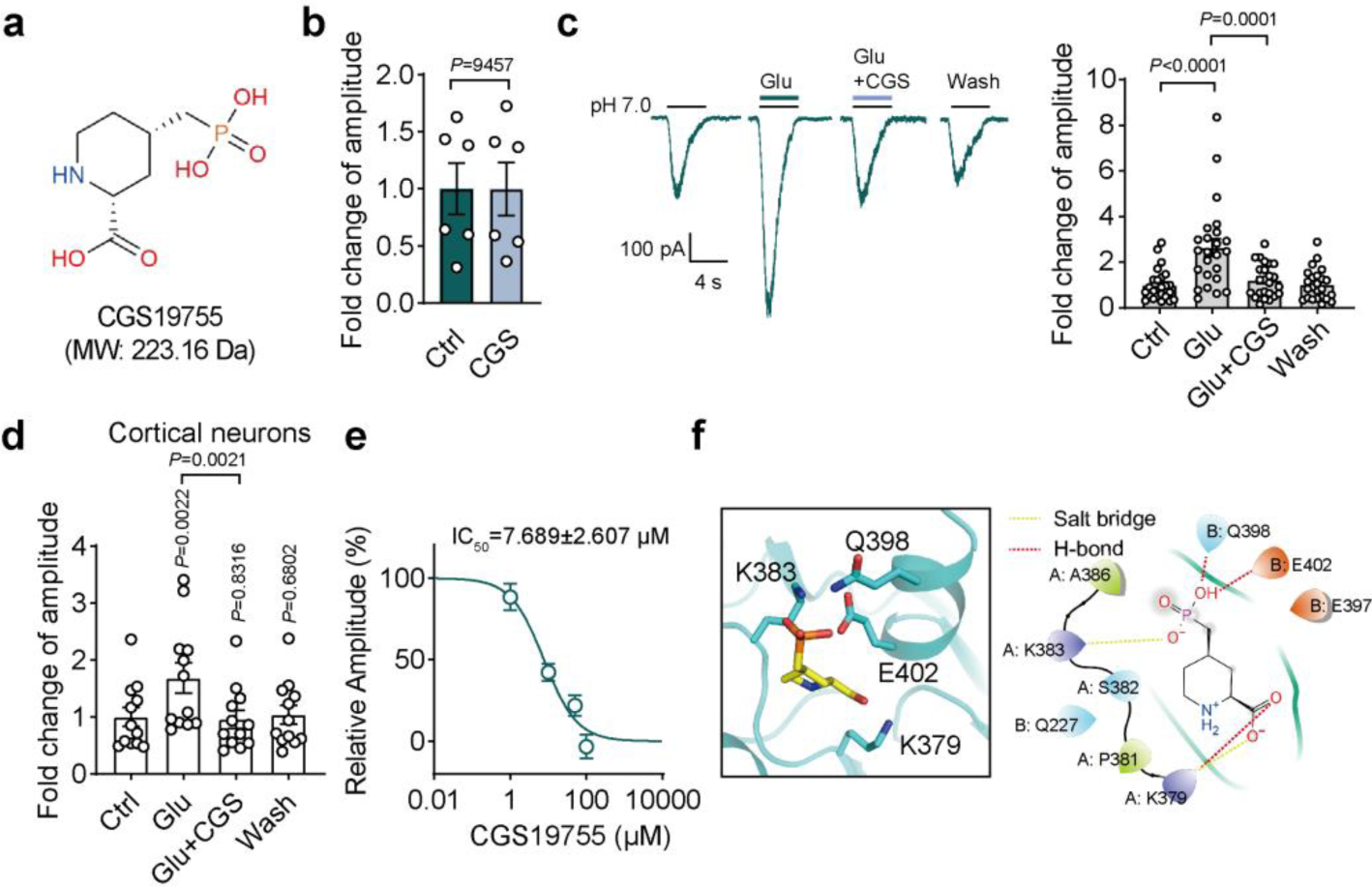
NMDAR antagonist CGS19755 blocks glutamate binding to ASIC1a. **a**, Chemical structure of CGS19755. **b**, 100 μM CGS19755, a competitive antagonists of NMDA receptors, had no effect on I_ASICs_. *n*=6 cells. **c**, **d**, CGS19755 can abolish glutamate-dependent potentiation of I_ASICs_ in ASIC1a transfected CHO cells (*n*=24 cells, **c**) and cultured cortical neurons (*n*=12 cells, **d**). **e**, Dose-response curve showing the percentage of inhibition of glutamate-dependent potentiation of I_ASICs_ by CGS19755. *n*=15 cells. **f**, Three-dimension and two-dimension images showing the glutamate binding pocket shared by CGS19755. The interactions between glutamate and surround residues are shown as yellow (salt bridge) and red (hydrogen-bond) dash lines. Data are mean±s.e.m.; two-tailed paired *t*-test (**b**); one-way ANOVA with Tukey post hoc correction (**c**,**d**). *P* values are indicated.

**Extended Data Figure 10.**
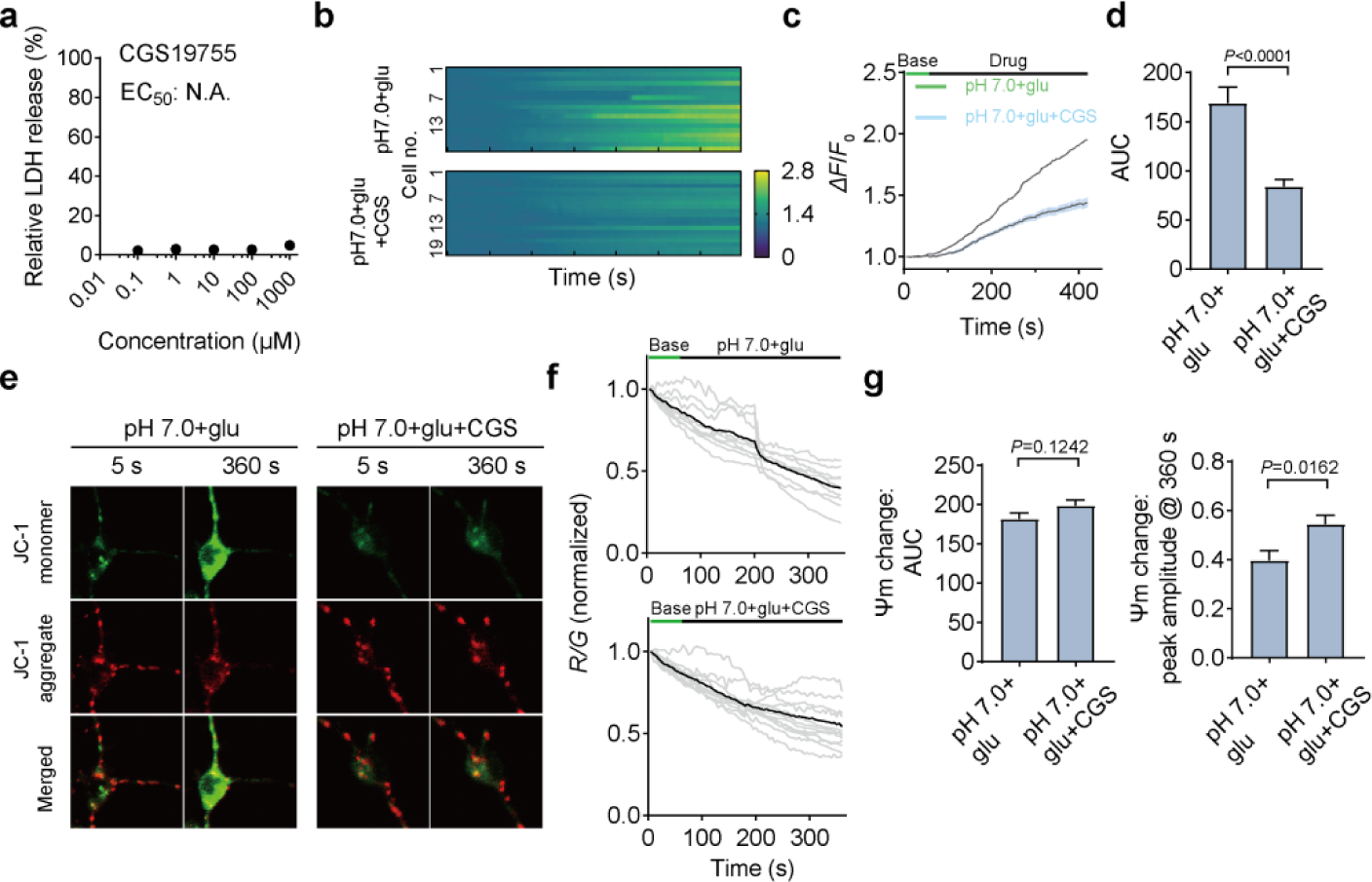
CGS19755 acts as a neuroprotectant against cell death *in vitro*. **a**, Determination of CGS19755 for basal cell death (1 hour of exposure to CGS19755) in primary cortical neurons by LDH release assay. *n*=5 wells for each concentration. N.A., not applicable. **b**,**c**, Calcium imaging of cultured cortical neurons recorded from wildtype mice with treatment of 500 μM glutamate alone or in combination with 100 μM CGS19755 in pH 7.0 solution. (**b**), fluorescence changes of individual cells; (**c**), average of fluorescence change of all cells from ***b***. **d**, Quantification of AUC from Ca^2+^ imaging under two treatment conditions. *n*=18 and 20 cells for each group. **e**,**f**, Representative images and traces showing changes of the mitochondrial potential of individual cells (grey) and their means (black) with different treatments of neurons from wildtype mice. When mitochondrial potential (absolute value) decreasing, fluorescence intensity of JC-1 monomer (green) enhanced, while JC-1 aggregate (red) faded. First frame at 5 second and last frame at 360 second are shown here (**f**). **g**, Quantifications of AUC of drug treatment (left panel) and peak amplitude (right panel) showing changes in mitochondrial membrane potential (Ψm) in pH 7 solution with glutamate alone or in combination with CGS19755. *n*=9 and 13 cells for each group. Data are mean±s.e.m.; two-tailed unpaired *t*-test (**d**,**g**). *P* values are indicated.

**Extended Data Figure 11.**
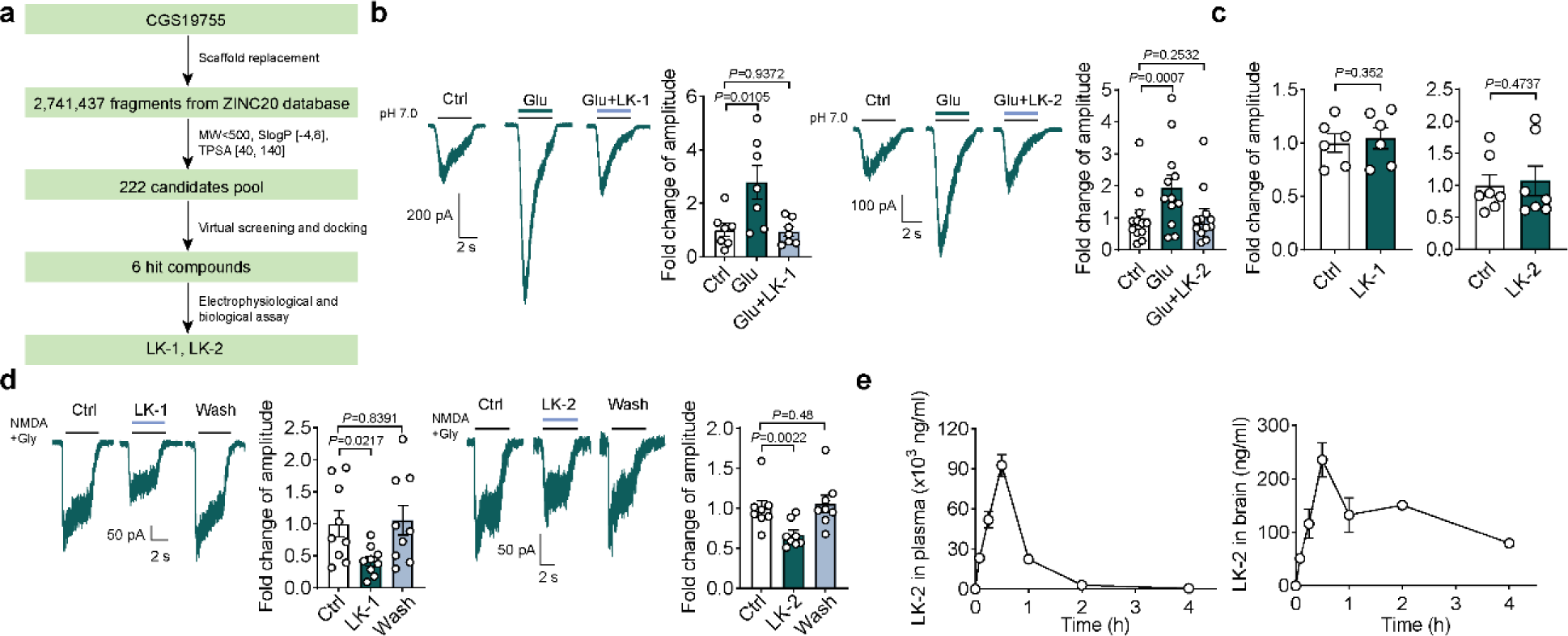
*In silico* screening identifies LK-2 over LK1 with more favorable pharmacological and pharmacokinetics profiles as a candidate for stroke therapy. **a**, Flow chart showing computational virtual screening of LK-1 and LK-2 based on the backbone structure of CGS19755. **b**, Glutamate-dependent enhancement of I_ASICs_ was inhibited by 100 μM LK-1 (*n*=7 cells) or 100 μM LK-2 (*n*=12 cells) in ASIC1a transfected CHO cells. **c**, 100 μM LK-1 (*n*=7 cells) or 100 μM LK-2 (*n*=12 cells) did not affect basal I_ASICs_ in ASIC1a transfected CHO cells. **d**, 100 μM LK-1 (*n*=9 cells) or 100 μM LK-2 (*n*=8 cells) inhibited NMDAR currents in acute isolated cortical neurons. 100 μM NMDA and 1 μM glycine were applied. **e**, Pharmacokinetic test *in vivo* showing the concentration of LK-2 in plasma and brain detected by LC-MS/MS. *n*=4 samples for each time point. Data are mean±s.e.m.; two-tailed paired *t*-test (**c**); one-way ANOVA with Tukey post hoc correction (**b**,**d**). *P* values are indicated.

**Extended Data Figure 12.**
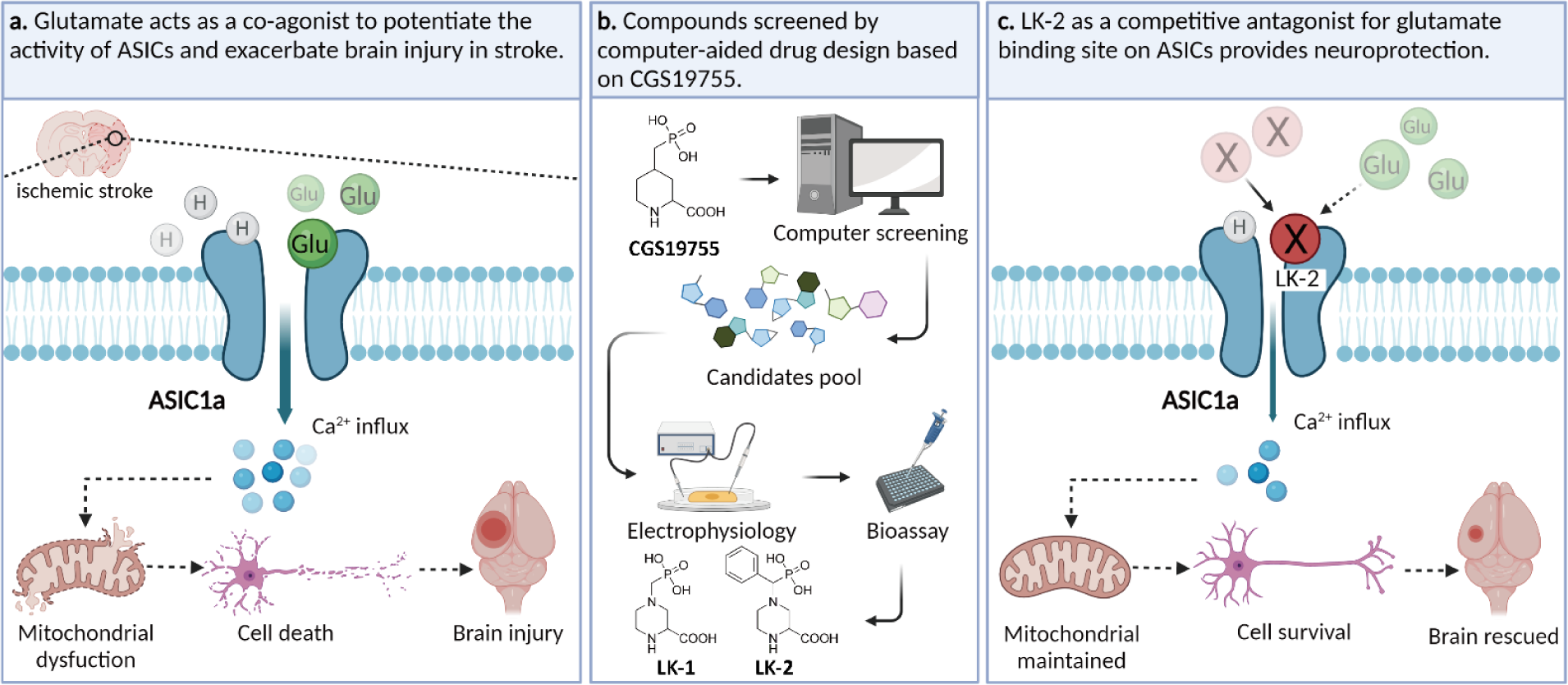
Diagram illustrating glutamate gates ASIC1a to mediate brain injury in mice stroke model. **a**, excessively released glutamate during stroke acts as a co-agonist and binds to ASIC1a, causing overactivation of ASICs and overload of Ca^2+^, triggering a cascade of cell death signaling including mitochondrial dysfunction and consequently brain injury. **b**, computational screening of NMDAR antagonist CGS19755 structural analogs leads to strong candidates, LK-1 and LK-2, being competitive antagonist for glutamate binding site on ASICs. **c**, LK-2 binding to ASIC1a precludes glutamate from binding to ASICs and attenuates the potentiating effect of glutamate without affecting the physiological gating of ASICs (and NMDARs), promoting cell survival and neuroprotection against ischemic brain injury. Diagram was assembled by BioRender.com.

**Extended data table 1.** Glutamate-cASIC1a binding site screening by HADDOCK.

|  | R161 | K379 | K383 | Q226 | K391 | K387 |
| --- | --- | --- | --- | --- | --- | --- |
| **HADDOCK score** | 5.6±0.3 | 8.3±5.9 | 17.4±8.8 | 19.5±5 | 22.1±5.4 | 22.8±3.3 |
| **E<sub>vdw</sub>** | -5.9±2 | -6.8±1.3 | -11.2±0.8 | -7.2±1 | -7.7±1.1 | -5.5±1.9 |
| **E<sub>elec</sub>** | -117.2±27.5 | -125.3±30.8 | -84.1±10.5 | -48.7±30 | -41.7±27.8 | -80.2±38 |
| **E<sub>desol</sub>** | 23.1±3.1 | 27.5±6.8 | 36.9±8 | 31.2±3.5 | 33.9±3.5 | 36.2±1.8 |
| **E<sub>air</sub>** | 1±0.27 | 0.5±0.21 | 1±0.72 | 0.4±0.18 | 0.5±0.12 | 0.6±0.12 |
| **BSA** | 287.1±12.3 | 336.2±7.6 | 353.1±12.3 | 313.7±14.7 | 253.3±37.4 | 226.8±23.7 |

The scoring is performed according to the weighted sum (HADDOCK score) of the following terms:
E<sub>vdw</sub>: van der Waals intermolecular energy;
E<sub>elec</sub>: electrostatic intermolecular energy;
E<sub>desol</sub>: desolvation energy;
E<sub>air</sub>: ambiguous interaction restraint energy;
BSA: buried surface area.

**Extended data table 2.** Impact of K379A mutation on stability of cASIC1a structure.

| Structure | Seq_length | pH | Mutation | Score | Delta_score | ddG | ddG_confidence |
| --- | --- | --- | --- | --- | --- | --- | --- |
| 5wku-mon-<br>omer chain<br>A | 417 | 6.9 | K379A | -7.459 | 0.013 | -0.089 | 0.897 |
| 5wku-trimer<br>chains ABC | 1249 | 6.9 | K379A | -13.127 | -0.165 | 0.003 | 0.886 |

The stability was evaluated by using MAESTRO [Laimer J. et al. (2015) MAESTRO-multi agent stability prediction upon point mutations. BMC Bioinformatics, 16, 116.], a structure-based method for predicting protein stability upon a mutation in a monomeric or multimeric form. The stability impact is summarized as a predicted free energy change (ddG) with prediction confidence given as ddG\_confidence.

**Extended data table 3.** CGS19755 binding pocket in cASIC1a.

|  |  |
| --- | --- |
| Overlap | 100% |
| Ligand volume | 190.98 Å<sup>3</sup> |
| Pocket volume | 783.76 Å<sup>3</sup> |
| Proportion of polar alpha spheres in the pocket | 0.89 (0~1) |
| Hydrophobicity score | 24.69 (0~100) |
| ALA | 2 |
| ASP | 2 |
| GLN | 1 |
| GLY | 1 |
| HIS | 2 |
| MET | 2 |
| PHE | 2 |
| SER | 1 |

**Extended data table 4.** LK-2 pharmacokinetic parameters in plasma and brain after a single intraperitoneal administration to male C57BL/6 mice.

| Parameter | i.p. 30 mg/kg |  |
| --- | --- | --- |
|  | Plasma | Brain |
| AUC<sub>(0-t)</sub> (µg/L*h) | 69793.793±3116.495 | 523.378±27.885 |
| AUC<sub>(0-∞)</sub> (µg/L*h) | 70278.227±3131.89 | 792.881±68.039 |
| MRT<sub>(0-t)</sub> (h) | 0.693±0.019 | 1.762±0.078 |
| MRT<sub>(0-∞)</sub> (h) | 0.721±0.018 | 3.63±0.611 |
| t<sub>1/2</sub> (h) | 0.576±0.015 | 2.297±0.441 |
| t<sub>max</sub> (h) | 0.5 | 0.5 |
| CL (L/h/kg) | 0.428±0.019 | 38.042±3.197 |
| V (L/kg) | 0.356±0.021 | 124.755±15.127 |
| C<sub>max</sub> (µg/L) | 92660±8312.196 | 235.875±31.904 |
| Brain/Plasma AUC ratio | 0.0127±0.0006 |  |

**Abbreviations:**
AUC: area under the concentration-time curve;
MRT: mean residence time;
t<sub>max</sub>: time to maximal concentration;
t<sub>1/2</sub>: terminal elimination half-life;
CL: clearance rate;
V: apparent volume of distribution;
C<sub>max</sub>: maximal concentration.

**Extended data table 5.** Physiological parameters of MCAO mice in Figure 2.

| Mice | Treatment | Body weight (g) | Body temperature (°C) | n |
| --- | --- | --- | --- | --- |
| WT | Sham | 19.8±0.18 | 32.9±0.15 | 3 |
|  | Saline | 19.9±0.15 | 31.2±0.11 | 11 |
|  | Memantine | 20.42±0.12 | 32.65±0.08 | 10 |
| *Asic1a*<sup>-/-</sup> | Saline | 22.5±0.19 | 33.17±0.1 | 9 |

**Physiological parameters of MCAO mice in Figure 4.**
| Mice | Treatment | Body weight (g) | Body temperature (°C) | n |
| --- | --- | --- | --- | --- |
| WT | Saline | 24.6±0.45 | 33.71±0.08 | 8 |
|  | CGS19755 | 27±0.25 | 33.28±0.3 | 12 |
|  | CGS19755+PcTX1 | 20.46±0.17 | 33.57±0.06 | 14 |
|  | LK-2 (10mg/kg) | 21.08±0.54 | 32.3±0.16 | 6 |
|  | LK-2 (30mg/kg) | 23.21±0.1 | 33.29±0.19 | 10 |
| *Asic1a*<sup>-/-</sup> | CGS19755 | 21.41±0.2 | 33.54±0.05 | 11 |

Data are mean ± s.e.m.

## References

1 Dirnagl, U., Iadecola, C. & Moskowitz, M. A. Pathobiology of ischaemic stroke: an integrated view. Trends in neurosciences 22, 391–397, doi:10.1016/s0166-2236(99)01401-0 (1999).

2 Lee, J. M., Zipfel, G. J. & Choi, D. W. The changing landscape of ischaemic brain injury mechanisms. Nature 399, A7–14, doi:10.1038/399a007 (1999).

3 O’Collins, V. E. et al. 1,026 experimental treatments in acute stroke. Ann Neurol 59, 467–477, doi:10.1002/ana.20741 (2006).

4 Xiong, Z. G. et al. Neuroprotection in ischemia: blocking calcium-permeable acidsensing ion channels. Cell 118, 687–698, doi:10.1016/j.cell.2004.08.026 (2004).

5 Gao, J. et al. Coupling between NMDA receptor and acid-sensing ion channel contributes to ischemic neuronal death. Neuron 48, 635–646, doi:10.1016/j.neuron.2005.10.011 (2005).

6 Yan, J., Bengtson, C. P., Buchthal, B., Hagenston, A. M. & Bading, H. Coupling of NMDA receptors and TRPM4 guides discovery of unconventional neuroprotectants. Science 370, doi:10.1126/science.aay3302 (2020).

7 Zong, P. et al. Functional coupling of TRPM2 and extrasynaptic NMDARs exacerbates excitotoxicity in ischemic brain injury. Neuron 110, 1944–1958 e1948, doi:10.1016/j.neuron.2022.03.021 (2022).

8 Collaborators, G. B. D. L. R. o. S., et al. Global, Regional, and Country-Specific Lifetime Risks of Stroke, 1990 and 2016. The New England journal of medicine 379, 2429–2437, doi:10.1056/NEJMoa1804492 (2018).

9 Benveniste, H., Drejer, J., Schousboe, A. & Diemer, N. H. Elevation of the extracellular concentrations of glutamate and aspartate in rat hippocampus during transient cerebral ischemia monitored by intracerebral microdialysis. Journal of neurochemistry 43, 1369–1374, doi:10.1111/j.1471-4159.1984.tb05396.x (1984).

10 Simon, R. P., Swan, J. H., Griffiths, T. & Meldrum, B. S. Blockade of N-methyl-D-aspartate receptors may protect against ischemic damage in the brain. Science 226, 850–852, doi:10.1126/science.6093256 (1984).

11 Park, C. K., Nehls, D. G., Graham, D. I., Teasdale, G. M. & McCulloch, J. The glutamate antagonist MK-801 reduces focal ischemic brain damage in the rat. Ann Neurol 24, 543–551, doi:10.1002/ana.410240411 (1988).

12 Keana, J. F. et al. Synthesis and characterization of a series of diarylguanidines that are noncompetitive N-methyl-D-aspartate receptor antagonists with neuroprotective properties. Proceedings of the National Academy of Sciences of the United States of America 86, 5631–5635, doi:10.1073/pnas.86.14.5631 (1989).

13 Grotta, J. C. et al. CGS-19755, a competitive NMDA receptor antagonist, reduces calcium-calmodulin binding and improves outcome after global cerebral ischemia. Ann Neurol 27, 612–619, doi:10.1002/ana.410270605 (1990).

14 Alvarez de la Rosa, D., et al. Distribution, subcellular localization and ontogeny of ASIC1 in the mammalian central nervous system. The Journal of physiology 546, 77–87, doi:10.1113/jphysiol.2002.030692 (2003).

15 Wemmie, J. A. et al. Acid-sensing ion channel 1 is localized in brain regions with high synaptic density and contributes to fear conditioning. The Journal of neuroscience : the official journal of the Society for Neuroscience 23, 5496–5502 (2003).

16 Chassagnon, I. R. et al. Potent neuroprotection after stroke afforded by a double-knot spider-venom peptide that inhibits acid-sensing ion channel 1a. Proceedings of the National Academy of Sciences of the United States of America 114, 3750–3755, doi:10.1073/pnas.1614728114 (2017).

17 Heusser, S. A. & Pless, S. A. Acid-sensing ion channels as potential therapeutic targets. Trends in Pharmacological Sciences, doi:10.1016/j.tips.2021.09.008 (2021).

18 Paukert, M., Babini, E., Pusch, M. & Grunder, S. Identification of the Ca2+ blocking site of acid-sensing ion channel (ASIC) 1: implications for channel gating. The Journal of general physiology 124, 383–394, doi:10.1085/jgp.200308973 (2004).

19 Grunder, S. & Pusch, M. Biophysical properties of acid-sensing ion channels (ASICs). Neuropharmacology 94, 9–18, doi:10.1016/j.neuropharm.2014.12.016 (2015).

20 Faden, A. I., Demediuk, P., Panter, S. S. & Vink, R. The role of excitatory amino acids and NMDA receptors in traumatic brain injury. Science 244, 798–800, doi:10.1126/science.2567056 (1989).

21 Schinder, A. F., Olson, E. C., Spitzer, N. C. & Montal, M. Mitochondrial dysfunction is a primary event in glutamate neurotoxicity. The Journal of neuroscience : the official journal of the Society for Neuroscience 16, 6125–6133 (1996).

22 White, R. J. & Reynolds, I. J. Mitochondrial depolarization in glutamate-stimulated neurons: an early signal specific to excitotoxin exposure. The Journal of neuroscience : the official journal of the Society for Neuroscience 16, 5688–5697 (1996).

23 Vergun, O., Keelan, J., Khodorov, B. I. & Duchen, M. R. Glutamate-induced mitochondrial depolarisation and perturbation of calcium homeostasis in cultured rat hippocampal neurones. The Journal of physiology 519 **Pt** 2, 451–466, doi:10.1111/j.1469-7793.1999.0451m.x (1999).

24 Yoder, N., Yoshioka, C. & Gouaux, E. Gating mechanisms of acid-sensing ion channels. Nature 555, 397–401, doi:10.1038/nature25782 (2018).

25 Sun, D. et al. Structural insights into human acid-sensing ion channel 1a inhibition by snake toxin mambalgin1. Elife 9, doi:10.7554/eLife.57096 (2020).

26 Laimer, J., Hofer, H., Fritz, M., Wegenkittl, S. & Lackner, P. MAESTRO--multi agent stability prediction upon point mutations. BMC Bioinformatics 16, 116, doi:10.1186/s12859-015-0548-6 (2015).

27 Spahi, S. et al. Hit identification against peptidyl-prolyl isomerase of Theileria annulata by combined virtual high-throughput screening and molecular dynamics simulation approach. Comput Biol Chem 89, 107398, doi:10.1016/j.compbiolchem.2020.107398 (2020).

28 Curtis, D. R. & Watkins, J. C. The excitation and depression of spinal neurones by structurally related amino acids. Journal of neurochemistry 6, 117–141, doi:10.1111/j.1471-4159.1960.tb13458.x (1960).

29 Lenda, F. et al. Synthesis of C5-tetrazole derivatives of 2-amino-adipic acid displaying NMDA glutamate receptor antagonism. Amino Acids 40, 913–922, doi:10.1007/s00726-010-0713-1 (2011).

30 Brauner-Osborne, H. et al. A new highly selective metabotropic excitatory amino acid agonist: 2-amino-4-(3-hydroxy-5-methylisoxazol-4-yl)butyric acid. Journal of medicinal chemistry 39, 3188–3194, doi:10.1021/jm9602569 (1996).

31 Brauner-Osborne, H., Nielsen, B. & Krogsgaard-Larsen, P. Molecular pharmacology of homologues of ibotenic acid at cloned metabotropic glutamic acid receptors. European journal of pharmacology 350, 311–316, doi:10.1016/s0014-2999(98)00246-5 (1998).

32 Perez-Pinzon, M. A. et al. Correlation of CGS 19755 neuroprotection against in vitro excitotoxicity and focal cerebral ischemia. J Cereb Blood Flow Metab 15, 865–876, doi:10.1038/jcbfm.1995.108 (1995).

33 Ge, Y., Chen, W., Axerio-Cilies, P. & Wang, Y. T. NMDARs in Cell Survival and Death: Implications in Stroke Pathogenesis and Treatment. Trends Mol Med 26, 533–551, doi:10.1016/j.molmed.2020.03.001 (2020).

34 Davis, S. M., Albers, G. W., Diener, H. C., Lees, K. R. & Norris, J. Termination of Acute Stroke Studies Involving Selfotel Treatment. ASSIST Steering Committed. Lancet 349, 32, doi:10.1016/s0140-6736(05)62166-6 (1997).

35 Grotta, J. et al. Safety and tolerability of the glutamate antagonist CGS 19755 (Selfotel) in patients with acute ischemic stroke. Results of a phase IIa randomized trial. Stroke; a journal of cerebral circulation 26, 602–605, doi:10.1161/01.str.26.4.602 (1995).

36 Sveinbjornsdottir, S. et al. The excitatory amino acid antagonist D-CPP-ene (SDZ EAA-494) in patients with epilepsy. Epilepsy research 16, 165–174, doi:10.1016/0920-1211(93)90031-2 (1993).

37 Investigators, T. N. A. G. A. i. N. G. Phase II studies of the glycine antagonist GV150526 in acute stroke : the North American experience. Stroke; a journal of cerebral circulation 31, 358–365, doi:10.1161/01.str.31.2.358 (2000).

38 Lees, K. R. et al. Tolerability of the low-affinity, use-dependent NMDA antagonist AR-R15896AR in stroke patients: a dose-ranging study. Stroke; a journal of cerebral circulation 32, 466–472, doi:10.1161/01.str.32.2.466 (2001).

39 Hardingham, G. E. & Bading, H. Synaptic versus extrasynaptic NMDA receptor signalling: implications for neurodegenerative disorders. Nature reviews. Neuroscience 11, 682–696, doi:10.1038/nrn2911 (2010).

40 Wang, Y. Z. et al. Tissue acidosis induces neuronal necroptosis via ASIC1a channel independent of its ionic conduction. Elife 4, doi:10.7554/eLife.05682 (2015).

41 Wang, J.-J. et al. Disruption of auto-inhibition underlies conformational signaling of ASIC1a to induce neuronal necroptosis. Nature Communications 11, doi:10.1038/s41467-019-13873-0 (2020).

42 Miesenbock, G., De Angelis, D. A. & Rothman, J. E. Visualizing secretion and synaptic transmission with pH-sensitive green fluorescent proteins. Nature 394, 192–195, doi:10.1038/28190 (1998).

43 Gonzalez-Inchauspe, C., Urbano, F. J., Di Guilmi, M. N. & Uchitel, O. D. Acid-Sensing Ion Channels Activated by Evoked Released Protons Modulate Synaptic Transmission at the Mouse Calyx of Held Synapse. The Journal of neuroscience : the official journal of the Society for Neuroscience 37, 2589–2599, doi:10.1523/JNEUROSCI.2566-16.2017 (2017).

44 Qi, X. et al. Pharmacological Validation of ASIC1a as a Druggable Target for Neuroprotection in Cerebral Ischemia Using an Intravenously Available Small Molecule Inhibitor. Frontiers in pharmacology 13, 849498, doi:10.3389/fphar.2022.849498 (2022).

45 Kreple, C. J. et al. Acid-sensing ion channels contribute to synaptic transmission and inhibit cocaine-evoked plasticity. Nature neuroscience 17, 1083–1091, doi:10.1038/nn.3750 (2014).

46 John A. Wemmie, J. C., Candice C. Askwith, Alesia M. Hruska-Hageman, Margaret P. Price, Brian C. Nolan, Patrick G. Yoder, Ejvis Lamani,1 Toshinori Hoshi, John H. Freeman, Jr., and Michael J. Welsh. The Acid-Activated Ion Channel ASIC Contributes to Synaptic Plasticity, Learning, and Memory. Neuron 34, 463–477 (2002).

47 Taugher, R. J. et al. The bed nucleus of the stria terminalis is critical for anxiety-related behavior evoked by CO2 and acidosis. The Journal of neuroscience : the official journal of the Society for Neuroscience 34, 10247–10255, doi:10.1523/JNEUROSCI.1680-14.2014 (2014).

48 Wang, Q. et al. Fear extinction requires ASIC1a-dependent regulation of hippocampalprefrontal correlates. Sci Adv 4, eaau3075, doi:10.1126/sciadv.aau3075 (2018).

49 Pignataro, G., Simon, R. P. & Xiong, Z. G. Prolonged activation of ASIC1a and the time window for neuroprotection in cerebral ischaemia. Brain : a journal of neurology 130, 151–158, doi:10.1093/brain/awl325 (2007).

## References

1 Li, H. et al. Alternative splicing of GluN1 gates glycine site-dependent nonionotropic signaling by NMDAR receptors. Proceedings of the National Academy of Sciences of the United States of America 118, doi:10.1073/pnas.2026411118 (2021).

2 Chu, X. P., Close, N., Saugstad, J. A. & Xiong, Z. G. ASIC1a-specific modulation of acidsensing ion channels in mouse cortical neurons by redox reagents. The Journal of neuroscience : the official journal of the Society for Neuroscience 26, 5329–5339, doi:10.1523/JNEURO-SCI.0938-06.2006 (2006).

3 Song, X. L. et al. Postsynaptic Targeting and Mobility of Membrane Surface-Localized hASIC1a. Neurosci Bull 37, 145–165, doi:10.1007/s12264-020-00581-9 (2021).

4 Li, M., Kratzer, E., Inoue, K., Simon, R. P. & Xiong, Z. G. Developmental change in the electrophysiological and pharmacological properties of acid-sensing ion channels in CNS neurons. The Journal of physiology 588, 3883–3900, doi:10.1113/jphysiol.2010.192922 (2010).

5 Spahi, S. et al. Hit identification against peptidyl-prolyl isomerase of Theileria annulata by combined virtual high-throughput screening and molecular dynamics simulation approach. Comput Biol Chem 89, 107398, doi:10.1016/j.compbiolchem.2020.107398 (2020).

6 Malde, A. K. et al. An Automated Force Field Topology Builder (ATB) and Repository: Version 1.0. J Chem Theory Comput 7, 4026–4037, doi:10.1021/ct200196m (2011).

